# The impact of pro-inflammatory cytokines on the β-cell regulatory landscape provides new insights into the genetics of type 1 diabetes

**DOI:** 10.1101/560193

**Authors:** M. Ramos-Rodríguez, H. Raurell-Vila, ML. Colli, MI. Alvelos, M. Subirana, J. Juan-Mateu, R. Norris, JV. Turatsinze, ES. Nakayasu, BJ. Webb-Robertson, JRJ. Inshaw, P. Marchetti, L. Piemonti, M. Esteller, JA. Todd, TO. Metz, DL. Eizirik, L. Pasquali

## Abstract

Early stages of type 1 diabetes (T1D) are characterized by local autoimmune inflammation and progressive loss of insulin-producing pancreatic β cells. We show here that exposure to pro-inflammatory cytokines unmasks a marked plasticity of the β-cell regulatory landscape. We expand the repertoire of human islet regulatory elements by mapping stimulus-responsive enhancers linked to changes in the β-cell transcriptome, proteome and 3D chromatin structure. Our data indicates that the β cell response to cytokines is mediated by the induction of novel regulatory regions as well as the activation of primed regulatory elements pre-bound by islet-specific transcription factors. We found that T1D-associated loci are enriched of the newly mapped cis-regulatory regions and identify T1D-associated variants disrupting cytokine-responsive enhancer activity in human β cells. Our study illustrates how β cells respond to a pro-inflammatory environment and implicate a role for stimulus-response islet enhancers in T1D.

---

In type 1 diabetes (T1D) early inflammation of the pancreatic islets (insulitis) by T and B cells contributes to both the primary induction and secondary amplification of the immune assault, with inflammatory mediators such as the cytokines interleukin-1β (IL-1β) and interferon-γ (IFN-γ) contributing to the functional suppression and apoptosis of β cells^1–3^.

Genome wide association studies (GWAS) have made a substantial contribution to the knowledge of T1D genetic architecture uncovering >60 regions containing thousands of associated genetic variants. Nevertheless, translating variants to function remains a main challenge for T1D and other complex diseases. Most of the associated variants do not reside in coding regions^4^ suggesting that they may influence transcript regulation rather than altering protein coding sequences. Recent studies showed a primary enrichment of T1D association signals in T and B cells enhancers^4,5^. A secondary^5^, or a lack of enrichment, was instead observed in islet regulatory regions. While such observation point to a major role of the immune system, we hypothesize that a subset of T1D variants may also act at the β-cell level but only manifest upon islet-cell perturbation and are not captured by the current maps of islet regulatory elements.

We have now mapped inflammation-induced *cis*-regulatory networks, transcripts, proteins and 3D chromatin structure changes in human β cells (**Fig. 1a**). We leverage these data to unmask functional T1D genetic variants as well as key candidate genes and regulatory pathways contributing to the β cell autoimmune destruction. Such analyses permit elucidation of the role of epigenetic gene regulation and its interaction with T1D genetics in the context of the autoimmune reaction that drives β cell death.

**Figure 1.**
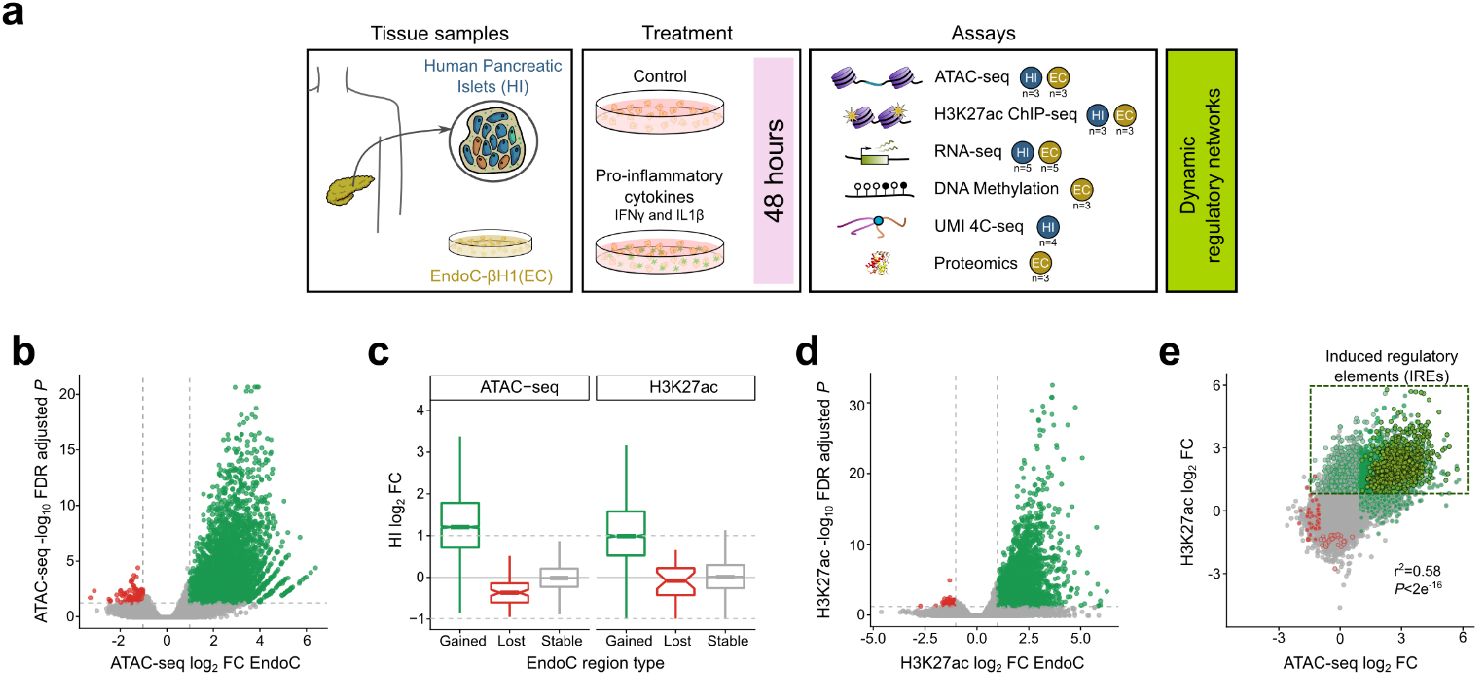
Pro-inflammatory cytokines exposure causes profound changes in the human pancreatic β-cells regulatory landscape. **a,** Summary of the experimental design. Multi-omics experiments were performed upon exposure or not of human pancreatic islets and EndoC-βH1 cells to IFN-γ and IL-1β to reconstruct cytokine-responsive dynamic regulatory networks. The number of human pancreatic islets (HI) and EndoC-βH1 (EC) replicate sample used in different assays is annotated at the side of each assay schematic representation. Note that we are currently extending our analysis to include additional replicate samples, **b, d,** Volcano plots of ATAC-seq **(b)** and H3K27ac ChlP-seq **(d)** changes obtained after exposure of EndoC-βH1 to IFN-γ and IL-1 β; green and red dots correspond to sites with |log_2_ FC|>1 and FDR adjusted *P*<0.05 as calculated by fitting a negative binomial model in DESeq2. Chromatin changes are classified as “gained” and “lost” chromatin sites whereas non-significant changes are defined as “stable”, **c,** Chromatin changes observed in EndoC-βH1 are largely replicated in human pancreatic islets exposed to the same treatment as illustrated by the distribution of log_2_ FC obtained from ATAC-seq and H3K27ac in human islet samples at open chromatin or H3K27ac enriched regions as classified in **b** and **d** in EndoC-βH1. Dotted lines indicate log_2_ FC thresholds (|log_2_ FC |>1). Box plot limits show upper and lower quartiles, whiskers extend to 1.5 times the interquartile range and the notch represents the confidence interval around the median, **e,** Correlation between chromatin accessibility and H3K27ac deposition. Each dot corresponds to a chromatin site, the x-axis indicating the ATAC-seq signal log_2_ FC and y-axis the log_2_ FC of the H3K27ac ChlP-seq signal. Point fill refers to the ATAC-seq and the border to the H3K27ac classification (gained=green; lost=red; stable=grey). The dotted box depicts the regulatory elements referred as induced regulatory elements (IREs) and the lighter shade of green depicts a subtype of these IREs named neo regulatory elements (see text).

## Results

### Pro-inflammatory cytokines have a profound impact on the pancreatic β-cell chromatin landscape

To characterize the effect of pro-inflammatory cytokines on the β-cell regulatory landscape we first mapped all accessible or open chromatin sites in human pancreatic islets exposed or not to IFN-γ and IL-1β. We assayed chromatin accessibility by ATAC-seq and, in order to focus on the β-cell fraction and to decrease inter-individual variability, in parallel with human pancreatic islet assays, we performed ATAC-seq in the clonal human β-cell line EndoC-βH1^6^, exposed or not to the pro-inflammatory cytokines (overall number of peaks identified in human islets: 91,600-195,015; and in EndoC-βH1 cells: 52,735-93,387. **Supplementary Fig. 1a** and **Supplementary Table 6**). Such experiments unmasked an important remodeling of the β-cell chromatin resulting in ~5,000 high confident chromatin sites that gain accessibility (FDR adjusted *P*<0.05; |log_2_ FC|>1) (**Fig. 1b**) upon exposure to pro-inflammatory cytokines. Importantly, the changes observed in the human β cell line were concordant with those observed in the human islet preparations (**Fig. 1c** and **Supplementary Fig. 1b**).

We reasoned that changes in chromatin accessibility may reflect the activation of non-coding *cis*-regulatory elements. We thus used chromatin immunoprecipitation coupled with next generation sequencing (ChIP-seq) to map cytokine-induced changes of H3K27ac (**Supplementary Fig. 1a-b**), a key histone modification associated with active *cis*-regulatory elements that was shown to be dynamically regulated in response to acute stimulation^7^. We observed genome-wide deposition of the active histone modification mark upon exposure to pro-inflammatory cytokines in both EndoC-βH1 and human pancreatic islets (**Fig. 1c-d**).

Integrative analysis of ATAC-seq and ChIP-seq indicates that changes in chromatin accessibility are strongly correlated with deposition of H3K27ac (*P*<2×10^-16^, r^2^=0.58) allowing the identification of ~2,600 open chromatin regions that gained H3K27ac (FDR adjusted *P*<0.05; |log_2_ FC|>1) upon exposure to pro-inflammatory cytokines (**Fig. 1e** and **Supplementary Fig. 1b**). We found that this subset of open chromatin regions is preferentially located distally to gene transcription start sites (TSS) (**Supplementary Fig. 1c**), their sequence is evolutionary conserved (**Supplementary Fig. 1d**) and enriched for specific transcription factor (TF) binding sites (**Supplementary Fig. 1e**). We named these newly mapped regions IREs for *“Induced Regulatory Elements’’* (**Supplementary Table 1**).

### Cytokine-induced regulatory elements drive β cell transcriptomic and proteomic changes

We next explored whether the newly identified IREs were associated with changes in gene expression and protein translation. To identify β-cell transcripts and proteins induced by the pro-inflammatory cytokines we assayed gene expression by RNA-seq (five replicates in EndoC-βH1 and five replicates in human pancreatic islets^8^, **Supplementary Fig. 1a**) and collected multiplex proteomics data for three EndoC-βH1 replicates after exposure or not to pro-inflammatory cytokines.

In line with the chromatin assays, that indicated extensive gene regulatory activation, we unraveled cytokine-induced transcriptional activation resulting in ~1,400 upregulated genes (FDR adjusted *P*<0.05; |log_2_ FC|>1) (**Supplementary Fig. 2a-b**). By multiplex proteomics, after rigorous filtering, a subset of 10,166 proteins was confidently quantified and retained for significance testing. A total of 348 proteins displayed significant changes in abundance (FDR/Q-value <0.15 and |FC|>1.5; |log_2_ FC|>0.58) being 2.19% of the overall detected proteins upregulated (**Supplementary Fig. 2c**), 76% of which had induced mRNA levels at 48h, confirming consistency between RNA-seq and protein changes (r^2^=0.72, *P*<2×10^-16^) (**Fig. 2a**). Protein-protein interactions inferred from β-cell cytokine-induced proteins resulted in a network more connected than expected by chance (*P* < 10^-3^), significantly enriched for Molecular Signatures Database (MSigDB; http://software.broadinstitute.org/gsea/msigdb/) pathways including IFN-γ signaling, antigen processing and presentation, apoptosis and T1D (KEGG T1D *P*=7.9×10^-8^, **Supplementary Fig. 2d**).

**Figure 2.**
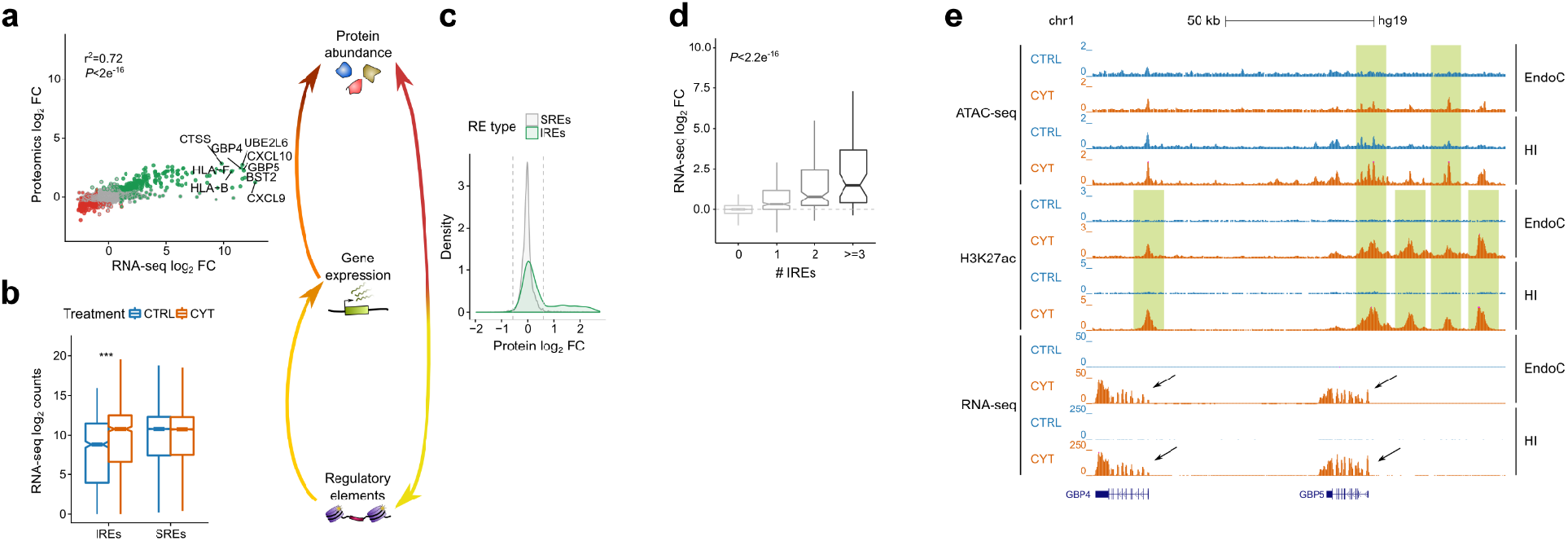
Cytokine-induced chromatin remodeling is coupled with changes in transcription and protein abundance. **a,** Correlation between changes in RNA expression and the corresponding protein abundance. The x-axis shows the log_2_ FC for the RNA expression and the y-axis the log_2_ FC for the protein abundance (both in EndoC-βH1 cells). Point fill indicates the RNA-seq classification and the border indicates the protein classification (up-regulated=green; down-regulated=red; equal-regulated=grey). **b,** Genes located <20 kb from IREs show cytokine-induced expression in EndoC-βH1 cells exposed or not to pro-inflammatory treatment. CYT= cytokine exposed (orange), CTRL = not exposed to cytokine (blue). *** Wilcoxon test *P*<0.001. **c,** Abundance of proteins encoded by IRE-associated genes is induced by cytokine exposure in EndoC-βH1 cells. This is shown by the significantly different (*P*<10^-16^) log_2_ FC distribution of protein abundance obtained after exposure or not of EndoC-βH1 to IFN-γ and IL-1β (n=3), for proteins encoded by genes located <20 kb from IREs or stable regulatory elements (SREs). **d,** An additive effect on gene up-regulation was observed for multiple IREs located at <20kb of a gene. Box plot limits show upper and lower quartiles, whiskers extend to 1.5 times the interquartile range and the notch represents the confidence interval around the median. ANOVA *P*<2.2×10^-16^. **e,** Representative view of the IFN-inducible guanylate binding proteins *GBP4* and *GBP5,* showing the up-regulation of these genes upon cytokine exposure and the nearby induction of IREs characterized by gains in chromatin accessibility and enrichment in H3K27ac (green boxes).

In line with our expectations we found that IREs were linked to up-regulation of the nearby gene/s as well as to an induced abundance of the corresponding protein (**Fig. 2b-c** and **Supplementary Fig. 2e**). Moreover, gene induction was highly correlated with the number of associated IREs, suggesting a cumulative effect of IREs on cytokine-induced changes in gene expression (**Fig. 2d**).”

Taken together these findings reveal that pancreatic β-cell response to pro-inflammatory cytokines is dynamic, involving extensive chromatin remodeling and profound changes in the regulatory landscape (**Fig. 2e** and **Supplementary Fig. 2f**). Such changes are associated with induction of transcription and protein translation including pathways implicated in the pathogenesis of T1D.

### The β cell response to pro-inflammatory cytokines is mediated by primed and neo regulatory elements

We next sought to gain insight into the dynamic activation of IREs. The relationship between chromatin openness and H3K27ac deposition upon exposure to pro-inflammatory cytokines allows the distinction of two classes of IREs (**Fig. 1e and Fig. 3a-c**): *“opening IREs’’* (n=1,198) which gain both chromatin accessibility (log_2_ FC>1) and H3K27ac (log_2_ FC>1); and *“primed IREs”* (n=1,455) which are already accessible chromatin sites prior to the treatment (ATAC-seq log_2_ FC<1) and gain H3K27ac (log_2_ FC>1) upon exposure to the stimulus. Primed and opening IREs are both associated to gene expression induction (**Supplementary Fig. 3a**), phylogenetically conserved (**Supplementary Fig. 3b**) and preferentially mapped distally relatively to gene TSS (**Supplementary Fig. 3c**). We further unmasked that 70% of opening IREs (n=838) are regions of complete novel activation (i.e. undetectable in basal condition, see methods). We named the latter *“neo IREs’’.* Neo IREs represent 30% of all IREs and may mirror *“latent enhancers*’’ that were identified upon stimulation of mouse macrophages^7^.

Because chromatin openness, the feature discerning the two classes of IREs, is believed to reflect TF occupancy, we analyzed their sequence composition in search of recognition sequences of key TFs orchestrating the β cell response to pro-inflammatory cytokines. Even though IREs are mostly distal to TSS (**Supplementary Fig. 3c**), in order to reduce sequence biases, we excluded from this analysis all annotated promoters, focusing exclusively on distal regulatory elements that displayed patterns of putative cytokine-responsive enhancers. The two classes of IREs showed clear differences in sequence composition. Newly induced enhancers were enriched for binding motifs of inflammatory-response TFs including Interferon-Sensitive Response Element (ISRE), STAT, IRF and NF-kB (**Supplementary Fig. 3d**). Primed enhancers instead were enriched for binding motifs of inflammatory-response TFs (ISRE, STAT, BARHL2), and unexpectedly, islet-specific TFs (HNF1A/B, NEUROD1, FOXA2, MAFA/B) (**Supplementary Fig. 3e**). Importantly, we found that, in primed enhancers, inflammatory-response and islet-specific TFs binding motifs mapped to the same genomic region, suggesting co-binding and possibly cooperation of the two classes of TFs (**Supplementary Fig. 3f-g**).

Sequence composition bias *per se* does not imply TF occupancy. We thus took advantage of published ChIP-seq datasets of islet-specific TFs (MAFB, PDX1, FOXA2, NKX6.1 and NKX2.2) mapped in un-stimulated human pancreatic islets^9^ to measure TF occupancy in primed and neo enhancers prior to the pro-inflammatory stimulus. As expected from the sequence composition analysis, primed enhancers (unlike neo enhancers) are highly bound by tissue-specific TFs even before their activation (**Fig. 3d and Supplementary Fig. 3h**). TF occupancy can also be indirectly assessed by ATAC-seq, which assays the protection of the bound sequence to transposase cleavage (footprint). Footprint analysis is effective for TFs with a long residence time^10^ such as IRFs and STAT transcription factor families. Our analyses revealed the emergence of footprint marks upon pro-inflammatory treatment in correspondence to ISRE motifs in both primed and neo enhancers (**Fig. 3e**) indicating cytokine-induced TF occupancy of IRE.

Gene regulation is orchestrated by different epigenetics mechanisms. DNA methylation is a relatively stable epigenetic mark contributing to maintenance of cellular identity^11,12^. Moreover, high-resolution DNA methylation maps, obtained from multiple tissues, established that the vast majority of tissue-specific differentially methylated regions are located at distal, mostly non-coding, regulatory sites^13^. Consequently, characterization of the DNA methylome in the context of relevant stimuli is important for understanding the functional mechanisms of tissue-specific responses in human disease^14^. We thus explored if cytokine-induced chromatin remodeling is associated with changes in DNA methylation. We quantified DNA methylation changes by performing dense methylation arrays in EndoC-βH1 exposed or not to IFN-γ and IL-1β. The Infinium MehtylationEPIC array was designed to interrogate with high precision and coverage >850,000 CpG sites (approximately 3% of all sites in the genome) selected primarily because of their location close to gene promoters and CpG-island regions. By focusing on the 962 IRE enhancers harboring one or more

CpG sites interrogated by the array, we observed that primed enhancers overlap lowly methylated CpGs (median β-value 0.16±0.14), that did not vary significantly upon cytokine exposure. Such observation is in sharp contrast with neo enhancers that were highly methylated under control condition (median β-value 0. 77±0.12) but underwent a significant loss of DNA methylation (Wilcoxon test, *P*=4.34e^-4^) upon the treatment. While we did not observe cytokine-induced methylation we found that 100 CpG probes were significantly demethylated (FDR adjusted *P*<0.05; β_cyt_-β_ctrl_≤-0.20) upon cytokine exposure (**Supplementary Fig. 3i**). Of all the genome-wide demethylated probes detected, 46% overlapped an IRE, 74% of which are located at neo enhancers (**Supplementary Fig. 3j**). These results suggest that neo enhancers are enriched for methylated CpGs that undergo preferential demethylation upon cytokine treatment whereas primed enhancers are enriched for unmethylated CpGs that do not change their methylation status upon the cytokine exposure (**Fig. 3f**).

Taken together these analyses lead to a model, in which pro-inflammatory cytokines elicit a regulatory response in β cells characterized by: 1) induction of novel distal regulatory elements coupled with reduction of DNA methylation and binding of inflammatory response TFs and 2) activation of regulatory elements pre-bound by islet-specific TFs and induced by inflammatory response TFs (**Fig. 3g**).

Collectively, these results allow reconstructing *cis*-regulatory networks activated in human pancreatic β cells upon exposure to the pro-inflammatory cytokines IFN-γ and IL-1β (**Supplementary Fig. 4a-c and Supplementary Table 2**).

### Activation of regulatory elements is coupled with changes in the 3D chromatin structure

Regulatory regions can exert control over genes at megabase distances through the formation of DNA loops. These loops are often confined within structures known as topologically associating domains (TADs)^15–17^. TADs are largely conserved upon evolution, are invariant in different cell types and have their boundaries defined by the regulatory scope of tissue-specific enhancers^18–20^. Our knowledge regarding the general characteristics and mechanisms of loops is improving^21,22^, but much less is known regarding mechanisms and functional significance of dynamic looping events during biological processes.

We took advantage of promoter capture Hi-C (pcHi-C) performed in human pancreatic islets^23^ to explore long-range interactions between gene promoters and cytokine-induced and invariant distant regulatory elements. Interestingly, we observed that the interaction confidence scores captured between IRE enhancers and gene promoters in untreated islets were significantly reduced compared with SREs enhancers (*P*=9×10^-5^) (**Supplementary Fig. 5a**). As this finding points to potential dynamic properties of the interaction maps, we next sought to investigate if cytokine-induced regulatory changes are linked to modification of the 3D chromatin structure and if induction of β-cell cytokine-responsive regulatory elements is coupled with the formation of novel DNA looping interactions.

Hi-C profiles are limited in sequencing coverage and library complexity resulting in maps of reduced resolution relative to regulatory maps of functional elements. On the other hand, 4C approaches are difficult to interpret quantitatively mainly due to potential amplification biases. We thus applied targeted chromosome capture with unique molecular identifiers (UMI-4C), a recently developed method^24^, to quantitatively measure interaction intensities in human islets before and after exposure to pro-inflammatory cytokines. We centered the conformation capture viewpoint to the promoter of 12 genes *(TNFSF10, GBP1, CIITA,* among others) whose expression was strongly induced by the cytokine exposure.

UMI-4C showed marked changes in the 3D chromatin structure at the analyzed loci. Promoters of the induced genes gained chromatin interactions with distal genomic regions reflecting the formation of new DNA looping events (**Fig. 4a-b and Supplementary Fig. 5b-d**). Importantly, such new contacts were preferentially engaged with newly mapped human islet cytokine-responsive IREs (**Fig. 4c**).

**Figure 3.**
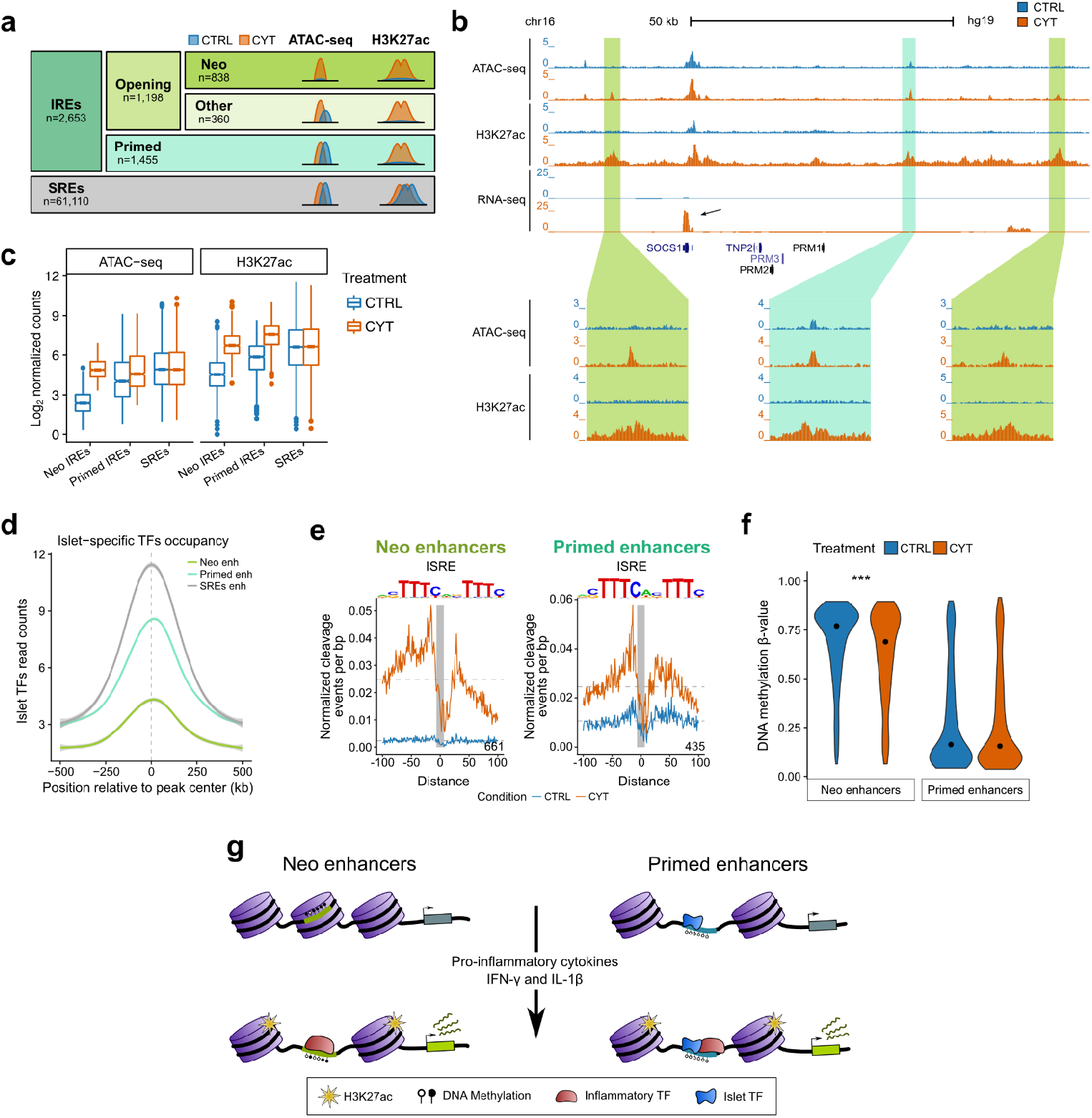
The β cell response to pro-inflammatory cytokine unveils neo and primed regulatory elements. **a,** Classification of ATAC-seq open chromatin sites upon exposure of human β cells to IFN-γ and IL-1β. IREs=induced regulatory elements, SREs=stable regulatory elements, **b**, View of the *SOCS1* locus, a gene strongly induced upon pro-inflammatory cytokine exposure. We here depict representative examples of primed (blue box) and neo IREs (green boxes), **c,** Box plot distribution of ATAC-seq and H3K27ac normalized tag counts at different classes of IREs. Box plot limits show upper and lower quartiles, whiskers extend to 1.5 times the interquartile range, individual data points represent outliers and the notch represents the confidence interval around the median. **d,** Islet-specific TF occupancy at neo, primed and stable regulatory elements. Read density for PDX1, NKX2.2, FOXA2, NKX6.1 and MAFB was calculated in 10bp bins in 1kb windows centered on the regulatory element. Lines represent means, while the grey shade depicts standard deviations. **e,** Footprint analysis of ISRE motifs in neo (left) and primed regulatory elements (right) in cells exposed or not to IFN-γ and IL-1β (control = blue; cytokines = orange). **f,** Violin plots showing the distribution of DNA methylation β-values in neo and primed enhancers, exposed or not to pro-inflammatory cytokines. *** *P*<0.001 Wilcoxon test. **g** Model showing two types of IREs driving the response to pro-inflammatory cytokines in human β cells.

**Figure 4.**
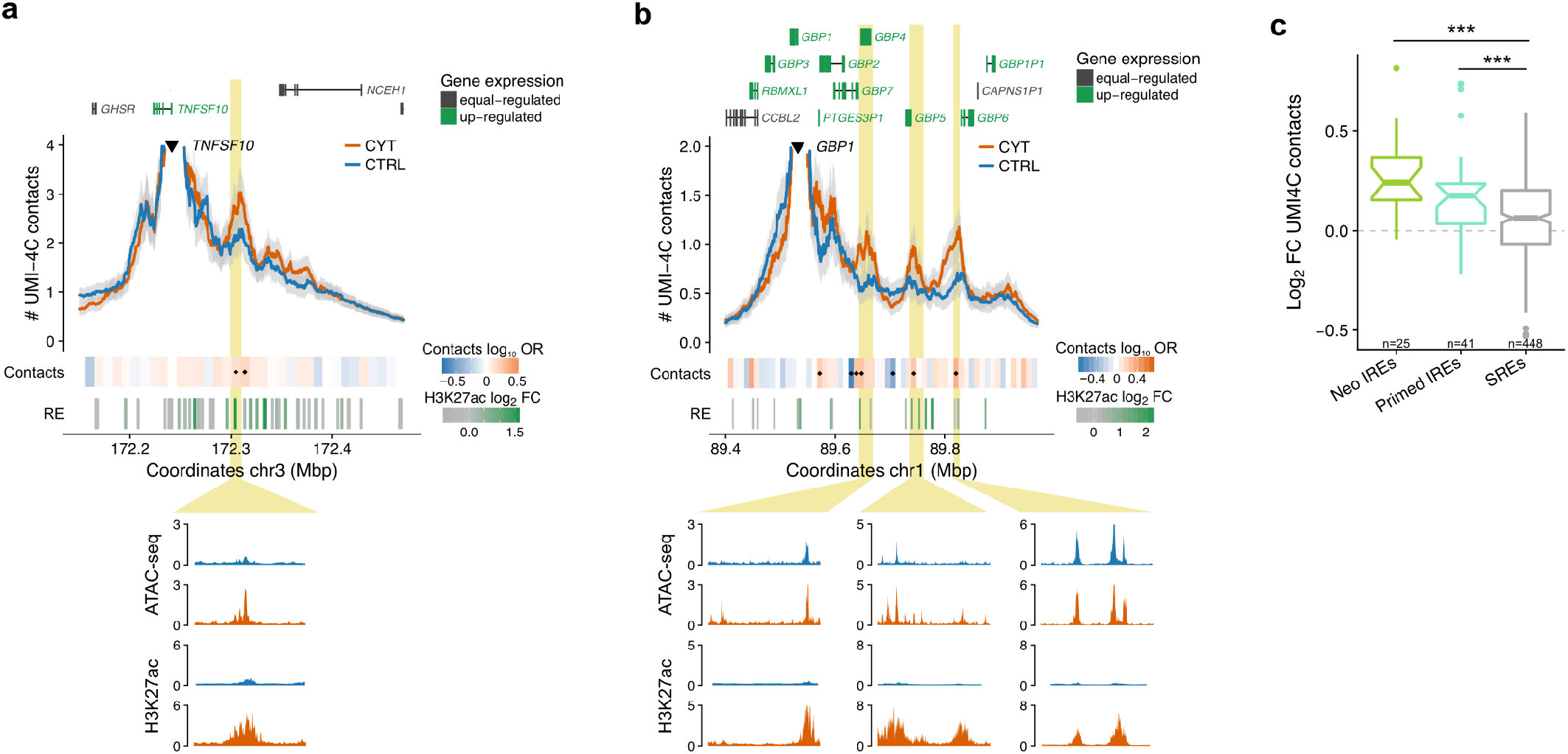
Cytokine exposure induces changes in human islet 3D chromatin structure. **a, b,** View of the UMI-4C chromatin contacts of *TNFSF10* **(a)** and *GBP1* **(b)** promoters, before and after exposure to pro-inflammatory cytokines. Yellow boxes indicate IREs that gain contacts with the up-regulated gene promoters. A heatmap under the 4C track represents the log_10_ odds ratio (OR) of the UMI-4C contacts difference in cytokine vs. control. Small black diamonds on top of the contact heatmap indicate a significant difference between cytokine-treated and control samples 3D chromatin contacts (*P*<0.05). ATAC-seq peaks are represented by rectangles shaded from gray to green proportionally to the cytokine-induced H2K27ac fold change observed at that site (RE=regulatory elements track). **c**, Distribution of the UMI-4C contacts fold changes (cytokines vs. control) at the different types of islets open chromatin sites classified as in **Fig. 3a**. The data obtained by analyzing 12 viewpoints centered at the promoter of cytokine-induced genes, show that the chromatin structural changes are preferentially happening at IREs. Box plot limits show upper and lower quartiles, whiskers extend to 1.5 times the interquartile range, individual data points represent outliers and the notch represents the confidence interval around the median.

These results demonstrate that cytokine exposure induces changes in human islet 3D chromatin conformation including the formation of novel enhancer-promoter interactions. Such changes allow the newly activated distal IREs to contact their target gene promoters.

### Cytokine induced islet regulatory elements are implicated in T1D genetic susceptibility

GWAS have identified more than 60 chromosome regions associated with T1D with many of the association signals having been assigned to candidate genes with immunological functions. Consistent with this notion, several studies reported a primary enrichment of T1D risk variants in T and B cell regulatory elements^4,5^. Furthermore, there is a substantial lack of statistical significant overlap of T1D associated variants in islet enhancers, while such regulatory elements are instead enriched for GWAS signals for T2D and fasting glucose^9,25^. Nonetheless, the molecular mechanisms linking T1D association signals to cellular functions remain poorly described for most of the regions of association identified.

We hypothesized that a subset of T1D genetic signals may reflect an altered capacity of the β cells to react to an inflammatory environment. We thus sought to explore to what extent genetic signals underlying T1D susceptibility act through pancreatic islet regulatory response to pro-inflammatory cytokines.

Causal *cis* variants are expected to lie in sequences that act as regulatory regions in state-specific and disease-relevant tissues. Consistent with this notion and, in line with previous observations^4,9^, by testing non shared loci associated with T2D and with T1D (the number of loci analyzed were 225 and 62 respectively), we found that T2D but not T1D risk loci overlap human islet non cytokine-responsive regulatory elements (i.e. SREs) more than expected by chance (SREs in T2D risk loci P= 2 x 10^-4^, Z=6.6). In contrast, we uncovered that T1D but not T2D risk loci are enriched for human islet IREs (IREs in T1D risk loci P=4×10^-3^, Z=4.8) (**Fig. 5a**). This result was reproduced when using regulatory elements detected in EndoC-βH1 cells (**Supplementary Fig. 6a**). Such findings unmasked 12 T1D associated regions (19% of the total) containing at least one islet IRE and including putative target genes whose expression was induced (**Supplementary Table 3**). All of these regions have the potential to harbor functional *cis*-regulatory variants, opening an avenue to the identification of T1D candidate genes acting at the β-cell level (**Supplementary Fig. 6b-f**).

**Figure 5.**
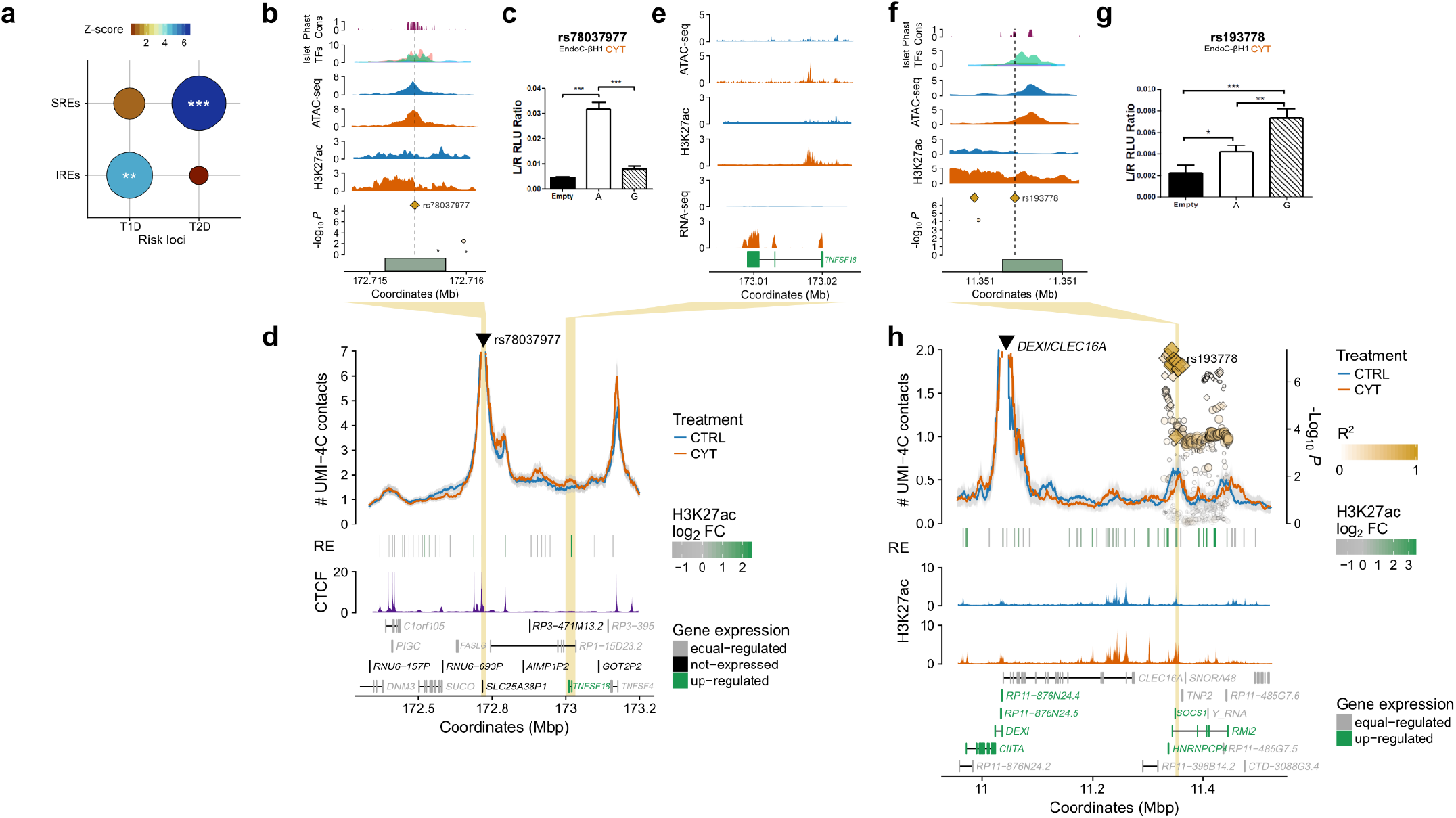
Cytokine-induced islet regulatory elements map to T1D associated regions and guide the identification of functional risk variants. **a**, T1D but not T2D risk loci are enriched for human islet IREs. In contrast T2D but not T1D risk loci overlap human islet stable regulatory elements (SREs) more than expected by chance. The permutation test results, assessing the significance of the overlap between different regulatory elements types (y axis) and trait-associated loci (x axis), are shown. The size of the circle is proportional to −log_10_ P; fill represents z-score of the observed versus expected value. Significance was assessed by permutation tests; * *P*<0.05, ** *P*<0.01, *** *P*<0.001. **b**, rs78037977 is the highest associated variant in the 1q24.3 T1D associated locus. The variant overlaps an IRE bound by islet-specific TFs under basal conditions. **c**, Luciferase assays in EndoC-βH1 exposed to pro-inflammatory cytokines show that, in these conditions, the sequence exerts enhancer activity which is reduced in the T1D associated allele (G) compared to the non-risk allele (A). **d**, UMI-4C analysis performed in human islets show that the cytokine-responsive regulatory element containing rs78037977 engages multiple distal chromatin contacts, some of which are induced upon INF-γ and IL-1β treatment and others coinciding with CTCFs binding sites. **e**, Zoom-in view at one induced chromatin contacts, mapping to an IRE in proximity of the up-regulated *TNFSF18* gene, a potential target of the IRE containing rs78037977 T1D functional variant. **f**, Variant rs193778 is part of the 99% credible set driving the association with T1D at the 16q 13.13 locus and maps to a phylogenetically conserved IRE. **g**, Luciferase assays for the underlying sequence performed in EndoC-βH1 exposed to INF-γ and IL-1β show significantly increased enhancer activity of the risk (G) allele compared to the non-risk (A) allele. **h**, UMI-4C analysis performed in pancreatic islets using the promoter of *DEXI* as viewpoint, show a chromatin contact with the IRE bearing the T1D variant. ATAC-seq peaks are represented, in d and h, by rectangles shaded from gray to green proportionally to the cytokine-induced H2K27ac fold change observed at that site (RE=regulatory elements track).

We noticed that the two T1D lead SNPs at 1q24.3 and 16q13.13 loci (rs78037977^26^ and rs193778^4^ respectively) were directly overlapping IREs in islets. We used GWAS genotyping data from a cohort of 14,575 individuals (5,909 T1D cases and 8,721 controls, see methods) to confirm their association with T1D. Both variants were included in the 99% credible set of their respective locus and displayed strong association *P*-values (rs78037977 *P*=6.94e^-10^; rs193778 *P*=1.33e^-7^; see Supplementary Table 4 for posterior probability of association and variant ranking in the credible set), indicating that they could be potentially causal.

At the 1q24.3 locus, rs78037977 overlaps an islet cytokine-induced chromatin site (**Fig. 5b**) which is pre-bound by islet-specific transcription factors and is a predicted enhancer in other cell types (**Supplementary Fig. 6g**). We created allele-specific luciferase reporter constructs and measured enhancer activity in the EndoC-βH1 line before and after cytokine exposure. The sequence exerts enhancer activity exclusively after cytokine exposure which is disrupted by the rs78037977, T1D associated, G allele (**Fig. 5c and Supplementary Fig. 6h**) consistent with a causal role of the variant at this locus. In order to identify the gene target of this T1 D-susceptible enhancer, we reconstructed the 3D chromatin structure by chromatin capture experiments. UMI-4C in human islets identified a cytokine-induced interaction of the enhancer with *TNFSF18* a gene activated in islets upon cytokine exposure (**Fig. 5d-e**). *TNFSF18* encodes for a cytokine, ligand of the TNFRSF18/GITR receptor, known to modulate inflammatory reaction and regulation of autoimmune responses^27^. Interestingly, we noticed that cytokine exposure results in upregulation of *TNFSF18* in human islets but not in the EndoC-βH1 β-cell line, suggesting differences in gene regulatory dynamics in primary tissue or the activation of an islet cell sub-population.

At the 16q13.13 locus, rs193778 maps to a phylogenetically conserved, cytokine-responsive regulatory element (**Fig. 5f and Supplementary Fig. 6g**). This sequence displays enhancer activity in both treated and untreated β cells. However, exclusively in cytokine-exposed β cells, the T1D-associated G allele exerts significantly higher enhancer activity than the protective variant (**Fig. 5g and Supplementary Fig. 6i**). The locus includes several up-regulated genes *(SOCS1, DEXI, CIITA, RMI2),* that could represent potential targets of this IRE. Recent works point to *DEXI* as a T1D candidate gene in immune cells and β cells^28,29^. By performing UMI-4C experiments in human islets we observed a strong chromatin contact between the promoter of *DEXI* and the regulatory element bearing the rs193778 T1D-associated variant (**Fig. 5h**), such data points to *DEXI* as a potential causal gene in pancreatic islets.

Altogether these results illustrate how unraveling cytokine-induced chromatin dynamics in human islets can guide the identification of *cis*-regulatory variants that are strong candidates in driving T1D-association signals.

## Discussion

Our work illustrates the human pancreatic β cell chromatin dynamics in response to an external stimulus that may be relevant in the context of T1D. We here show that exposure to pro-inflammatory cytokines causes profound remodeling of the β-cell regulatory landscape coupled with changes in gene expression and protein production. We are presently **replicating our experiments in additional human pancreatic islet samples** to confirm the consistence of our findings. We unveil the activation of ~2,600 cytokine-responsive distal *cis* regulatory elements and reveal a lack of homogeneity in their molecular mechanism of activation. We observed that the induction of a subset of novel regulatory regions (neo IREs) require TF binding and chromatin opening while other chromatin sites are “primed” to their activation being prebound by islet-specific TFs. Our observations suggest a model in which binding of tissue-specific TFs may facilitate chromatin accessibility at a subset of chromatin sites that can then be promptly activated by the induction of inflammatory-response TFs. Such model is supported by very recent findings^30^ and consistent with observations in murine macrophages^7,31^ and murine dendritic cells^32^, but thus far had not been demonstrated in a highly differentiated and non-immune related tissue such as the pancreatic islets.

Importantly, we show that such regulatory changes are coupled with 3D chromatin remodeling, allowing the newly activated regulatory elements to contact their target genes. Several reports described 3D chromatin dynamics properties in the cell developmental context^33,34^, upon loss of cell fate^35,36^, senescence^37,38,39^ or in response to hormonal exposure^39^. Our observations indicate that the capacity of enhancer loop formation is maintained in a highly-differentiated tissue such as the islets and it is coupled with transcriptional regulatory changes, in response to an external stimulus.

The model used in our study to explore chromatin dynamics is of particular interest because it mimics the inflammatory environment that the pancreatic islets may face in the early stages of T1D. While several T1D candidate genes regulating key steps related to “danger signal recognition” and innate immunity were shown to be expressed in human islets^40^, T1D associated variants were shown to be enriched for immune cell types but not in stable pancreatic islets regulatory elements^4^. Such apparent contradiction may be reconciled by our findings showing that T1D risk regions are enriched for human islet cytokine-responsive regulatory elements. Our data, supported by recent findings unmasking regulatory variants affecting enhancer activation in immune response^30,41^, opens the avenue to identify T1D molecular mechanisms acting at the pancreatic islet cells level.

Although we cannot exclude that functional variants disrupting β-cell regulatory mechanisms may at the same time affect the regulatory potential of immune-related cell types, the availability of stimulus-responsive cis-regulatory maps in pancreatic islets will facilitate hypothesis-driven experiments to uncover how common and lower frequency genetic variants impact islet cells in T1D. We here studied the islet response to a specific pro-inflammatory stimulus. Future work studying additional immune-mediated stresses potentially affecting β cells at different stages of the disease, may allow uncovering other association signals acting at the islet cell level.

In general, our findings could apply by extension to other diseases where “primed” enhancers may facilitate cell-type-specific responses to ubiquitous signals resulting in tissue-specific genetic susceptibility in autoimmune diseases.

## Acknowledgments

This work by LP was supported by grants from the Spanish Ministry of Economy and Competiveness (BFU2014-58150-R and SAF2017-86242-R to LP), Marató TV3 (201624.10, to LP) and a young investigator award from the Spanish Society of Diabetes to LP. The Pasquali lab is further supported by ISCIII (PIE16/00011). LP is a recipient of a Ramon y Cajal contract from the Spanish Ministry of Economy and Competitiveness (RYC-2013-12864) and MR is supported by an FI Agència de Gestió d’Ajuts Universitaris i de Recerca (AGAUR) PhD fellowship. JJ was supported by a MSCA fellowship grant from the Horizons 2020 EU program (Project reference: 660449). Human islets were provided through the European Consortium for Islet Transplantation (ECIT) distribution program for basic research supported by JDRF award 31-2008-416. DLE was supported by grants from the Fonds National de la Recherche Scientifique (FNRS), Welbio CR-2015A-06, Belgium; the Horizon 2020 Program, T2Dsystems (GA667191); Brussels Capital Region-Innoviris (project Diatype), and the Innovative Medicines Initiative 2 Joint Undertaking under grant agreement No 115797 (INNODIA). This Joint Undertaking receives support from the Union’s Horizon 2020 research and innovation programme and “EFPIA”, ‘JDRF” and “The Leona M. and Harry B. Helmsley Charitable Trust. TOM and DLE were supported by a grant of the National Institutes of Health, NIH-NIDDK-HIRN Consortium 1UC4DK104166-01. Part of the work was performed in the Environmental Molecular Sciences Laboratory, a U.S. Department of Energy (DOE) national scientific user facility at Pacific Northwest National Laboratory (PNNL) in Richland, WA. Battelle operates PNNL for the DOE under contract DE-AC05-76RLO01830. We thank Josep Mercader and Marta Guindo Martínez for helpful discussions regarding GWAS enrichment analyses.

## Author contributions

L.P., M.R. and D.L.E. designed the experiments. H.R., M.C., M.A., J.J., R.N. and E.N. performed and analyzed the experiments. M.R. performed bioinformatic analyses with contribution from M.S., J.T., B.W. and J.I. L.Pi., P.M., M.E., and T.M. provided material and resources. L.P., D.L.E., T.M. and J.T. supervised the study. LP and MR coordinated, conceived the project and wrote the manuscript with contribution from D.L.E. All authors reviewed the final manuscript.

## Data availability

Datasets for induced regulatory elements (IREs) in both EndoC-βH1 and human pancreatic islets will be available for download and visualization upon publication at the Islet Regulome Browser^42^ (www.isletregulome.com).

Raw sequencing reads for the different assays (RNA-seq, proteomics, ATAC-seq, ChIP-seq, UMI-4C) will be made available at GEO.

## Supplementary Materials

Supplementary tables

Supplementary Table 1. List of induced regulatory elements in EndoC-βH1 cells.

Supplementary Table 2. Annotation of induced regulatory elements to up-regulated genes.

Supplementary Table 3. T1D risk loci that overlap induced regulatory elements.

Supplementary Table 4. Credible sets for 1q24.3 and 16q13.13 loci.

Supplementary Table 5. Metadata from human islet samples.

Supplementary Table 6. Information about UMI-4C selected baits.

Supplementary Table 7. Oligonucleotides used for luciferase assays.

Supplementary Table 8. Sequencing statistics for the analyzed samples.

Supplementary Table 9. UMI-4C sequencing statistics.

Extended data

Extended Data 1. Bed file for called ATAC-seq peaks in EndoC-βH1.

Extended Data 2. Bed file for called ATAC-seq peaks in human islets.

Extended Data 3. Bed file for called H3K27ac ChIP-seq peaks in EndoC-βH1.

Extended Data 4. Bed file for called H3K27ac ChIP-seq peaks in human islets.

## Methods

### Ethics

Human islets were isolated from brain-dead organ donors in accordance with national laws and institutional ethical requirements at the Istituto Scientifico San Raffaele, Milan, Italy. All experiments were performed according to protocols approved by the institutional research committee of the Institute for Health Science Research Germans Trias i Pujol, Badalona, Spain.

### Human Islets

Pancreatic islets were isolated according to established isolation procedures^1,2^. After isolation, islets were incubated at 37°C in CMRL 1066 medium with 10% fetal calf serum prior to shipment at room temperature in the same culture medium. Upon arrival, samples were recultured at 37°C in Ham’s F-10 medium containing 10% fetal bovine serum (FBS), 2 mM GlutaMAX, 50 U/ml penicillin and 50 μg/ml streptomycin (GIBCO), 6.1 mM glucose, 50 μM 3-isobutyl-1-methylxanthine, 1% BSA (Sigma) for 1-3 days. The islets were then exposed or not to cytokines in the same medium without FBS for 48 hours. The following cytokine concentrations were used, as from previous dose-response experiments^3–5^: recombinant human IL-1β (specific activity 1.8×10^7^ U/mg; 201-LB-005, R&D Systems, Abingdon, UK) at 50 U/ml; recombinant human IFN-γ (specific activity 2×10^7^ U/mg; AF-300-02, Peprotech, London, UK) at 1000 U/ml as described^3^. Islets were next rinsed with phosphate buffered saline at room temperature, formaldehyde was added to a final concentration of 1%, and samples were incubated at room temperature with constant shaking for 10 minutes. Glycine was added to a final concentration of 125 mM for 5 minutes, and cells were rinsed two times with phosphate-buffered saline containing 1x protease inhibitor cocktail (Millipore) at 4°C. The cells were spun down at 1000 rpm for 4 minutes, snap frozen, stored at −80°C and then used for ChIP, and 4C experiments. Additional aliquots of the same sample were processed similarly but without exposing to formaldehyde or glycine and frozen for RNA extraction or directly processed for ATAC-seq experiments. Islet purity was initially assessed by dithizone staining using an aliquot of islets immediately prior to fixation. Only islet preparations with minimal exocrine contamination (purity >80%) were selected for the experiments described here (**Supplementary Table 5**). The glucose stimulation index^6^ (comparing 16.7 vs 3.3 mM glucose) on the preparations used ranged from 3.3 to 7.9, indicating that the islets were functionally competent. Cell viability was assessed in 4 of the 5 human islets exposed or not to IL-1β + IFN-γ and used for RNAseq, as previously described^7^. The results confirmed, as previously observed^7,8^, that the cytokine cocktail used in this study affects islet cell viability (control human islets viability 94 ± 1%; cytokine-exposed human islet viability 72 ± 4%. Paired T-test *P* = 0.02).

### EndoC-βH1

The human insulin-producing EndoC-βH1 cells, kindly provided by Dr R. Scharfmann, University of Paris, France^9^, were cultured in DMEM medium containing 5.6 mmol/l glucose, 2% BSA fraction V, 50 μmol/l 2-mercaptoethanol (Sigma-Aldrich, Poole, UK), 10 mmol/l nicotinamide (Calbiochem, Darmstadt, Germany), 5.5 μg/ml transferrin, 6.7 ng/ml sodium selenite (Sigma-Aldrich), 100 units/ml penicillin and 100 μg/ml streptomycin (Lonza, Leiden, Netherlands). The same concentrations of cytokines as described for the human islets experiments were used in the treatment of EndoC-βH1 cells^8^.

### ATAC-seq

ATAC-seq library preparations were carried out as previously described^10^ with minor modifications^11,12^. Briefly, we selected 50 healthy and acinar-free islets corresponding approximately to 50,000 cells and isolated their nuclei by incubating the islets in 300 μl cold lysis buffer for 25 min on ice and resuspending them after 5 and 15 min using a syringe with a 29G needle. The pellet was next washed in 100 μl of lysis buffer and centrifuged for 15 min at 500 *g* at 4°C. The transposition reaction was carried out in a 25 μl reaction mix containing 2 μl of Tn5 transposase, 12.5 μl of TD buffer (Nextera DNA Library Prep Kit, 15028212, Illumina, San Diego, USA) and 10.5 μl DEPC-treated water. The transposition reaction mix was incubated at 37°C for 1 h following inactivation by incubating for 30 min at 40°C after addition of 5 μl of clean up buffer (900 mM NaCl, 300 mM EDTA), 2 μl of 5% SDS and 2 μl of Proteinase K. Isolation of the tagmented DNA was performed with 2x SPRI beads cleanup (Agencourt AMPure XP, Beckman Coulter) and was eluted in 20 μl DEPC treated water.

Two sequential 9-cycle PCR were performed in order to enrich for small tagmented DNA fragments. The PCR mix consisted of 2 μl of PCR Primer 1 (25 μM working stock), 2 μl of Barcoded PCR Primer 2 (25 μM working stock; sequences provided in^10^), 25 μl of NeBNext High-Fidelity 2x PCR Master Mix, 1 μl of DEPC-treated water and 20 μl of the eluted sample (DEPC water was added to compensate in case the volume of eluted DNA was less than 20 μl). The library was amplified in a thermocycler using the following program (leave preheated to 72°C): 72 °C for 5 min; 98 °C for 30 s; 9 cycles of 98 °C for 10 s, 63 °C for 30 s; and 72 °C for 1 min; and at 4 °C hold. After the first PCR round, fragments smaller than 600bp were selected using SPRI cleanup beads. The DNA library was finally purified using the MinElute PCR Purification Kit (Qiagen, Hilden, Germany), following kit instructions and eluting 2 x 10 μl with the elution buffer. TapeStation was performed to check library quality and nucleosomal pattern of the fragments distribution resulted from the tagmentation reaction (**Supplementary Fig. 7a**). Semi-quantitative PCR assays at target positive and negative control sites were performed to estimate the efficiency of the ATAC-seq experiment before sequencing (data not shown).

### ChIP-seq

ChIP-seq was carried out using tagmentation (ChIPmentation) as previously described^13^. Briefly, 4,000 islet equivalents or ~4 million 1% formaldehyde fixed cells were lysed in sonication buffer (2% Triton X-100, 100mM NaCl, 10 mM Tris-HCl pH 8.0, 1 mM EDTA pH 8.0, 1% SDS, 1 × protease inhibitors cocktail (Millipore)) and sonicated with a Diagnode Bioruptor with the following conditions: 20 cycles of 1 minute (40 seconds on, 20 seconds off) at high power, to obtain a fragment size in the range of 100-500 bp. Lysates were centrifuged at 13,000 RPM for 5 min at 4°C, and the supernatant containing the sonicated chromatin was diluted with a ChIP dilution buffer (50mM Hepes pH 8.0, 1 mM EDTA pH 8.0, 140 mM NaCl, 0.75% Triton X-100, 0.1% sodium deoxycholate, 1 × protease inhibitors cocktail (Millipore)) to a final volume of 1ml per immunoprecipitation (IP). The antibody against H3K27ac epitope (1.5 μg per IP, Abcam ab4729) was added along with 50 μl 10% BSA and the IP was incubated over night at 4 °C. For each IP, 20 μl magnetic Protein A+G (Millipore) were washed twice and resuspended in PBS. The beads were then added to the IP and incubated for 2h at 4°C on a rotator. Blocked antibody-conjugated beads were then placed on a magnet and the supernatant was removed. Beads were subsequently washed with a low salt wash buffer (20 mM Tris-HCl pH 8.0, 2mM EDTA pH 8.0, 150 mM NaCl, 1% Triton X-100, 0.1% SDS), high salt wash buffer (20 mM Tris-HCl pH 8.0, 2mM EDTA pH 8.0, 500 mM NaCl, 1% Triton X-100, 0.1% SDS) and LiCl wash buffer (10 mM Tris-HCl pH 8.0, 1 mM EDTA pH 8.0, 250 mM LiCl, 1% DOC and 1% NP40). The IP was incubated for 4 minutes at 4°C with rotation for each wash buffer. Beads were then washed with cold 10mM Tris-Cl pH 8.0, to remove detergent, salts and EDTA. The whole reaction, including beads, was then transferred to a new tube and placed on a magnet to remove supernatant to decrease background. Beads were then carefully resuspended in 30 μl of the tagmentation reaction mix (10 mM Tris-HCl pH 8.0, 5 mM MgCl_2_) containing 1 μl Tn5 transposase from the Nextera DNA Library Prep Kit (15028212, Illumina, San Diego, USA) and incubated at 37°C for 10min. The beads were then washed twice with RIPA (10 mM Tris-HCl, pH 8.0, 1 mM EDTA, pH 8.0, 140 mM NaCl, 1% Triton x-100, 0.1% SDS, 0.1% DOC), once with cold Tris-EDTA. Beads were next incubated with 150 μl elution buffer (1% SDS, 0.1M NaHCO_3_) for 15 min with rotation at room temperature. The chromatin immunoprecipitate was separated from the beads using a magnet. The ChIP was then re-eluted by adding another 150 μl elution buffer to the beads. Both ChIP elutions were combined in the same tube. Each ChIP elute was then incubated with 0.75 μl RNaseA (20mg/ml) for 30 min at 37°C. 4.5 μl Proteinase K (Thermo Scientific), and 12 μl 5M NaCl was then added and the ChIP was incubated at 65°C overnight to revert formaldehyde cross-linking. Finally, ChIP DNA was purified using a Phenol-chloroform extraction protocol.

Two μl of each library were amplified in a 10 μl qPCR reaction containing 0.15 μM primers, 1 × SYBR Green and 5μl *NEBNext High-Fidelity 2X PCR Master Mix* (NEB M0541S), to estimate the optimum number of enrichment cycles with the following program: 72°C for 5 min, 98°C for 30 s, 24 cycles of 98°C for 10 s, 63°C for 30 s and 72°C for 30 s, and a final elongation at 72°C for 1 min. Final enrichment of the libraries was performed in a 50μl reaction using 0.75 μM primers and 25 μl *NEBNext High-Fidelity 2X PCR Master Mix.* Libraries were amplified for N+1 cycles, where N is equal to the rounded-up Cq value determined in the qPCR reaction. Enriched libraries were purified using SPRI AMPure XP beads at a beads-to-sample ratio of 0.7:1, followed by a size selection using AMPure XP beads to recover libraries with a fragment length of 100-500 bp. Semi-quantitative PCR assays at target positive and negative control sites were performed to estimate the efficiency of the ChIP experiment before sequencing (data not shown).

### RNA-seq

Total RNA was isolated from 400,000 EndoC-βH1 cells using the RNeasy Mini Kit (Qiagen) which retrieves RNA molecules longer than 200 nucleotides, as previously described in detail^14^. RNA integrity number (RIN) values were evaluated using the Agilent bioanalyzer 2100 (Agilent Technologies, Wokingham, UK). All the samples had RIN values of > 8. RNA-seq libraries generation was performed as described by the manufacturer (Illumina, Eindhoven, The Netherlands). In brief, mRNA was purified from 1-2 μg of total RNA using oligo (dT) beads, before it was fragmented and randomly primed for reverse transcription followed by second-strand synthesis to create fragments of double-stranded cDNA. The obtained cDNA had underwent paired-end repair to produce blunted ends. After 3’-monoadenylation and adaptor ligation, cDNAs were purified on an agarose gel and 200 bp products were excised from the gel. The purified cDNA was amplified by PCR using primers specific for the ligated adaptors. Finally, the libraries were submitted to quality control with the Agilent bioanalyzer 2100.

### Proteomics

For the proteomic analysis 1.5 million EndoC-βH1 cells treated or not with cytokines (IL-1β + IFN-γ) were processed using the Metabolite, Protein and Lipid Extraction (MPLEx) approach^15^. Protein pellets were dissolved in 50 mM NH**4**HCO**3** containing 8 M urea and 10 mM dithiothreitol and shaken at 800 rpm for 1 h at 37 °C. Sulfydryl groups were alkylated by adding 400 mM iodoacetamide (40 mM final concentration), and incubated for another hour in the dark at room temperature. Samples were then 8-fold diluted with 50 mM NH_4_HCO_3_, and CaCl_2_ was added to a final concentration of 1 mM from a 1 M stock solution. The digestion was carried out by adding trypsin at 1:50 enzyme:protein ratio and incubation at 37 °C for 5 h. The resulting peptides were extracted using C18 cartridges (Discovery, 50 mg, Sulpelco) and concentrated in a vacuum centrifuge. Peptides were then quantified by BCA, normalized and labeled with tandem mass tags (TMT-10plex, ThermoFisher Scientific) according to the manufacturer’s instructions. Labeled peptides were extracted using C18 cartridges and fractionated into 24 fractions using high-pH reversed phase chromatography^16^. Peptide fractions were loaded into a C18 column (70 cm × 75 μm i.d. packed with Phenomenex Jupiter, 3 μm particle size, 300 Å pore size) connected to a Waters NanoAquity UPLC system. A gradient of water (solvent A) and acetonitrile (solvent B) both containing 0.1% formic acid (1-8% B in 2 min, 8-12% B in 18 min, 12-30% B in 55 min, 30-45% B in 22 min, 45-95% B in 3 min, hold for 5 min in 95% B and 99-1% B in 10 min) was used to elute the peptides, which were directly analyzed by nanoelectrospray ionization on a Q-Exactive mass spectrometer (Thermo Fisher Scientific). Scans were collected with a resolution of 35,000 at 400 m/z in a 400-2000 m/z range. High-energy collision-induced dissociation (HCD) fragmentation were set for the 12 most intense parent ions using the following parameters: peptide charge ≥ 2, 2.0 m/z isolation width, 30% normalized collision energy and 17,500 resolution at 400 m/z. Each parent ion was fragmented only once before being dynamically excluded for 30 s.

Collected data were processed using Decon2LS_V2^17^ and DTARefinery^18^, both using default parameters, to recalibrate the runs and generate peak lists. Peptide identifications were done using MSGF+^19^ by searching peak lists against islet protein sequences deduced from a transcriptomics experiment^3^ and supplemented with keratin sequences (32,780 total protein sequences). For MSGF+ searches, a parent ion mass tolerance of 10 ppm, partially tryptic digestion and 2 missed cleavages were allowed. The following modifications were also considered during searches: cysteine carbamidomethylation and N-terminal/lysine TMT addition as static modifications, and methionine oxidation as a variable modification. Results were filtered in two steps to a final false-discovery rate <1%: spectral-peptide matches – MSGF probability ≤ 1.0E −9, and protein level < 1.0E −10. The intensity of TMT reporter ions was extracted using MASIC^20^.

Data quality of the multiple omics sets was assessed by evaluating of the distribution of the data for each sample via the robust Mahalanobis distance abundance vector (rMd-PAV) algorithm^21^. Data were then converted into log_2_ and normalized by standard median centering. Proteins were quantified using a Bayesian proteoform discovery methodology (BP-Quant) in combination with standard reference-based median quantification^22^ and were considered significant with a cutoff of p ≤ 0.05 based on a paired T-test.

Protein-protein interaction (PPI) network analysis was performed with GeNets^23^ using Metanetworks v1.0 that integrates PPI from InWeb3^24^ and ConsensusPathDB^25^. Default parameters were applied and Molecular Signatures Database (MSigDB; http://software.broadinstitute.org/gsea/msigdb/)^26^ enriched pathways were overlaid.

### UMI-4C data generation

UMI-4C was performed as described^27^ with minor modifications. Briefly, formaldehyde crosslinked human islets (estimated ~4 million individual cells) were treated with ice-cold lysis buffer (50 mM Tris-HCl pH 8, 150 mM NaCl, 1% TX-100, 0.5% IGEPAL-CA-630, Sigma; 5mM EDTA, AppliChem; 1X protease inhibitor cocktail, Millipore) and permeabilized with SDS (Millipore) and TX-100. Nuclei were digested with DpnII (New England Biolabs) for 16h at 37°C. Afterwards, more DpnII was added for 4h and ligated with T4 DNA ligase (Promega) at 16°C overnight. Cross-link was reversed by overnight incubation with Proteinase K at 65°C. Next, after 30 min incubation with 30 μl of 10 mg/ml RNase A at 37°C, DNA was purified by phenol/chloroform and ethanol precipitation. DNA was then resuspended in 10 mM T ris-HCl pH 8 and 5 ug aliquots were sonicated by Covaris S2 to fragment sizes in the range of 450-550 bp. Sonicated DNA was next incubated for 30 min at 20°C with 20 μl 10× end-repair buffer and 10 μl end-repair mix (NEB E6050S). The reaction was next cleared by 2× AmpureXP beads (Agencourt AMPure XP, Beckman Coulter) and eluted in 75 μl EB (10 mM Tris-HCl pH 8). A-tailing was performed on the elute by adding 10 μl NEB buffer 2, 4 μl Klenow fragment (NEB M0212L) and 10 μl 10 nM dATP 20 min at 37 °C and 20 min at 75°C to inactivate the reaction. DNA was next dephosphorylated with 2 μl calf intestinal alkaline phosphatase (NEB M0290S) and incubated at 50°C for 60 min and immediately purified by 2x AmpureXP beads. Illumina forked adaptors were added (final concentration 0.4 μM) and incubated with 80 μl quick ligase buffer and 5 μl of Quick Ligase (NEB M2200S) for 20 min at 25°C. DNA was next denatured at 96°C for 5 min, moved to ice for 5 min and then purified by 0,8x AmpureXP beads to release the non-ligated strand of the adaptor. Denatured DNA was quantified by Qubit (ssDNA HS Assay, Q32851, Invitrogen).

200 ng of DNA were used for library preparation by nested PCR. The first PCR reaction was performed with 10 μl GoTaq Flexi Buffer, 3 μl MgCl_2_ 25mM, 1 μl dNTP 10 mM, 2 μl US bait primer and 2 μl Illumina universal primer 10 μM (0,4 μM final concentration each primer) and 1 μl GoTaq (Promega, M5005) with the following program: 2 min 95°C, 20 cycles of 30 s 95°C, 30 s 56°C and 60 s 72°C and final extension of 5 min 72°C. After the first PCR reaction, the DNA was purified with 1× AmpureXP beads, and 31 μl of the elute was used to perform a second PCR reaction following the same conditions as the first one but using the DS bait primer and 17 amplification cycles instead of 20. Finally, DNA was purified by 0,8x AmpureXP beads. To increase molecular complexity, each library was obtained by pooling 5-10 PCRs per viewpoint. The PCR primers used in UMI-4C are listed in **Supplementary Table 6**. Each library was sequenced to a depth of >1M 75bp long paired-end reads using either NextSeq or HiSeq 2500 platforms.

### Luciferase reporter assays

For episomal reporter assays in the EndoC-βH1 cell line, selected human cytokine-induced regulatory elements regions were first amplified from genomic DNA with primers (**Supplementary Table 7**) containing XhoI/HindIII restriction sites. The amplicons were next cloned into the PGL4.23[luc2/minP] Luciferase Reporter Vector (Promega) as previously described^28^. Briefly, the amplicon and the vector were simultaneously digested. Next, the vector was dephosphorylated with FastAP (Thermo Scientific). The DNA was then purified and ligated with a T4 DNA ligase (Promega). Next, the generated Reporter Vectors were transformed into *E. coli* (DH5α) and purified using the Nucleospin Plasmid (740588.250, MN, Düren, Germany).

Site-directed mutagenesis was used to introduce single nucleotide variants into the generated construct. The variants were generated by PCR using the primers shown in **Supplementary Table 7**. The parental supercoiled double-stranded DNA was digested with *DpnI* (NEB, 174R0176S) 1h at 37°C and the constructs were transformed in competent *E. coli* cells (DH5a) by thermal shock. Finally, the introduced variants were checked by Sanger sequencing. EndoC-βH1 cells were transfected in 24 well plates, at a density of 300,000 cells per well, with 200 ng of reporter vectors or empty vectors, plus 20 ng of phRL-CMV Renilla-luciferase to control for transfection efficiency.

Transfections were performed with lipofectamine 2000 (Invitrogen) for 8h, according to manufacturer instructions. Upon transfection EndoC-βH1 medium was supplemented with 2% FBS^8^ and exposed or not to the cytokines for 48 h. After 48 h, the cells were assayed using the Dual Luciferase Assay (Promega, Madison, USA), following manufacturer instructions. The luciferase units were measured using VICTOR Multilabel Plate Reader (PerkinElmer). Firefly luciferase activity was normalized to Renilla luciferase activity and then divided by values obtained for the empty pGL4.23. The assays were performed in at least three independent experiments.

### ATAC-seq, ChIP-seq read mapping and data processing

ATAC-seq and ChIP-seq libraries were sequenced on Illumina HiSeq 2500 platform to generate 49-50 bp single or paired end reads. 42-180M ATAC-seq reads and 15-75M ChIP-seq reads were aligned to the hg19 reference genome using Bowtie 2 (version 2.3.4.1)^29^ with default parameters. After alignment, reads mapping to ENCODE blacklist regions^30^, to non-autosomal chromosomes or to mitochondrial DNA were discarded. Duplicates were removed using Picard Mark Duplicates (version 2.5.0)^31^ (see **Supplementary Table 8** for number of mapped reads per experiment and **Supplementary Fig. 7b-e** for quality control plots for ATAC-seq data). For ATAC-seq samples we performed an offset correction of 4 bp on the + strand and 5 bp on the – strand to adjust the read start sites to the center reads on the transposon’s binding event as previously described^10^ using an in-house script.

Open chromatin peaks were called with MACS2 (version 2.1)^32^ callpeak using the following parameters “-q 0.05 --nomodel --shift −100 --extsize 200”. H3K27ac enriched regions were identified with the same software using the following parameters “--broad --broad-cutoff 0.1 --nomodel”. We compared the number of called peaks with other published ATAC-seq datasets in human islets and beta bells^33–35^ and observed that >80% of the stronger peaks found in such datasets are also present in our merged peak set.

For both assays peaks were called separately for each replicate as a measure of quality control. Merged BAM files for each condition and experiment were converted to bedgraph using bedtools (version 2.26)^36^ genomeCoverageBed, RPKM scaled and transformed into bigWig (bedGraphToBigWig UCSC tool^37^) to be uploaded to a public server for visualization.

### RNA-seq read mapping and data processing

RNA-seq libraries were sequenced on a HiSeq 2000 plataform to produce 100bp long paired-end reads with an average of 180 million reads per replicate (EndoC-βH1; n=5).

Reads were aligned using TopHat (version 2.0.13)^38^ to GChr37 genome with default parameters. Afterwards, reads were assigned to Gencode version 18 gene annotation^39^ using htseq-count (version 0.6.1p1)^40^ with default parameters. The RNA-seq of 5 human islet preparations^41^ was used for comparison and processed in an identical way.

### Differential analysis of ATAC-seq, RNA-seq, ChIP-seq

For both ATAC-seq and ChIP-seq, aligned reads from all replicates and conditions were merged into a single BAM file to identify a comprehensive set of peaks. We next used such peak set to compute read counts, separately for each replicate and condition. In the case of RNA-seq data, the output of htseq-count^40^ was used as the input matrix for downstream analysis. The generated matrices were normalized and differential analysis was performed using DESeq2^42^ with default parameters. Briefly, DESeq2 algorithm performs size factors estimation, to normalize each sample by its library size; then, calculates dispersions for each feature to account for intra-sample variability; finally, it fits a negative binomial generalized linear model (GLM) and calculates the Wald statistic. To improve statistical power, DESeq2 performs an independent step to filter out low mean normalized counts are filtered out that bear a limited chance of being significant. To control type I error, raw p-values are adjusted using the Benjamini-Hochberg method to calculate a False Discovery Rate (FDR). Thresholds for significance were set at a FDR adjusted *P* =<0.05 and |log_2_ FC|≥1. All regions/genes that did not reach significance or did not pass log_2_ FC cutoff were classified as stable/equal-regulated.

### 850K Infinium MethylationEPIC Array data generation

DNA from EndoC-βH1 cells exposed or not to IL-1β and IFN-γ for 48h as described above (3 replicates per condition) was extracted using QIAamp DNA Mini kit (Qiagen, Venlo, The Netherlands). 1 μg DNA aliquots (n=3) were processed for 850K Infinium MethylationEPIC Array (Illumina) as previously described^43^. Briefly, DNA was bisulfite converted using EZ-96 DNA Methylation kit (Zymo Research Corp., CA, USA) and then hybridized as described in the Infinium Methylation Assay Protocol^44^.

The resulting array signals were processed and analyzed using RnBeads R package^45^ with the following functions: *rnb.run.qc,* for quality control, *rnb.run.preprocessing* for filtering and normalization, *rnb.run.differential* for performing the differential analysis, comparing samples exposed or not to pro-inflammatory cytokines. The method used for assessing differences between groups consists on fitting a hierarchal linear model as implemented in limma package^46^ using M-values (log of β-values) as a metrics to measure methylation levels^47^. CpGs were considered as differentially methylated when FDR adjusted *P*<0.05 and absolute difference in methylation β-values between cytokine and control samples was >0.2 (20% changes in methylation).

### Defining classes of induced regulatory elements

In order to characterize chromatin accessibility dynamics upon exposure of human islets and EndoC-βH1 to pro-inflammatory cytokines, we processed the results obtained from the DESeq2 differential analysis and computed the overlap between ATAC-seq peaks and H3K27ac enriched sites. Regions annotated as “stable” for both ATAC-seq and H3K27ac assays were classified as *Stable Regulatory Elements* (SREs). Regions classified as either “stable” or “gained” in ATAC-seq differential analysis and as “gained” in H3K27ac were classified as *Induced Regulatory Elements* (IREs).

*Induced Regulatory Elements* (IREs) were classified in two groups: “Opening” IREs (n=1,198), corresponding to regions annotated as “gained” for both ATAC-seq and H3K27ac and “primed” IREs (n=1,455) for regions annotated as “stable” for ATAC-seq and “gained” for H3K27ac. Since opening IREs include a gradient of cytokine-induced chromatin accessibility changes, we next selected only those opening regions that were completely closed prior to the cytokine exposure. For this purpose, we considered newly open chromatin those opening ATAC-seq peaks that were not called in the control samples using a relaxed threshold (*P*≤0.001). Such analysis allowed us to identify a subset of 838 opening regions that we named “neo” IREs. A similar approach was used to identify macrophage latent enhancers^48^. Finally, IREs identified in EndoC-βH1 that showed an H3K27ac FC >1.5 in human pancreatic islets were considered IREs consistently induced in both cell types (**Supplementary Fig. 1b**).

### Sequence conservation analysis

Sequence conservation at different classes of induced regulatory elements was assessed by determining average phastCons 46 way score in placental mammals^49^. The scores average was calculated in 50bp bins over a 2kb window center on the open chromatin site. A set of regions identical in number and size shuffled over the mappable genome using regioneR^50^ were used as control set.

### Assignment of regulatory elements to target genes

In order to annotate regulatory elements as distal or proximal, we assigned each regulatory element to the nearest TSS of a coding gene (using gencode v18 annotation^39^). Those regions that lie within 2kbs from the nearest TSS were annotated as promoters. The rest of the regulatory elements were considered enhancers.

To test association between different classes of open chromatin and changes in gene expression and protein abundance (**Fig. 2b-d, Supplementary Fig. 2e, 3a**) in an unbiased manner, we assigned ATAC-seq sites to a gene when closer than 20kb of its TSS.

Finally, in order to detect all possible IRE gene targets, we assigned to each IRE all up-regulated genes whose TSS was closer than 40 Kb. When an up-regulated gene could not be found in <40 Kb, the IRE was assigned to the closest, but <1Mb far, induced gene (**Supplementary Fig. 4a and Supplementary Table 2**).

### Sequence composition and transcription factor analysis

*De novo* motif analysis was performed using HOMER (version 4.8.2)^51^ *findMotifGenome.pl* tool with parameters “-size given -bits”. Only enriched sequences present in more than 1.5% of targets were retained. Selection of best matches was performed as follows: all matches with scores over 0.80 were included in the table. For those hits without any match over 0. 80, the first hit was selected and the score was included in the table.

To assay motif colocalization, we used all motifs instances identified in the *de novo* analysis in primed enhancers. First we used the *annotatePeaks.pl* tool from HOMER to map all these motif instances in primed enhancers and SRE enhancers. Next, the motif colocalization was calculated by counting motif pairs found in each ATAC-seq peak. Significance was determined by Fisher Exact test comparing colocalization of motifs pairs in IREs vs. SREs. Only significant pairs (*P*<0.001) were retained.

To evaluate islet-specific transcription factors occupancy, we used ChIP-seq bam files for PDX1, NKX2.2, FOXA2, NKX6.1 and MAFB^52^. We computed the read coverage in the regions of interest over 10bp bins. Reads were quantile-normalized, mean counts in each bin for each transcription factor were calculated and the mean for all TFs was plotted.

To identify footprints from the ATAC-seq data, we generated tag directories with all ATAC-seq replicates in each condition using HOMER *makeTagDirectory.* Neo and primed enhancers were centered on the ISRE motif matrix annotated with *annotatePeaks.pl* with options “-center motif1.motif-size given” and tag means for 5’ and 3’ read ends were obtained using *annotatePeaks.pl* with options “-size −100, 100 -hist 1 -d tagsDir”. The resulting 5’ ends were plotted using ggplot2^53^.

In order to create a non-redundant dataset of motifs for the gene regulatory analysis (**Supplementary Fig. 4a**), motifs from primed and opening enhancers were compared using the *compareMotifs.pl* script from HOMER. The motifs were then mapped to primed and opening enhancers using *annotatePeaks.pl.*

### Gene regulatory network analysis

Regulatory networks were constructed using 1.9 *networkx* python module^54^. IRE enhancers, harboring *de novo* enriched TF binding motifs, were linked to their inferred target gene as described above. A random network with the same number of nodes as the observed network and an edge creation probability p=0.1 was generated for comparison using the *erdos_renyi_graph* module. The inferred networks were visualized using Cytoscape (version 3.6.0)^55^ applying a force directed layout.

Functional enrichment analysis of the IREs gene targets (**Supplementary Fig. 4c**) was performed by Metascape^56^. The enrichment analysis for Gene Ontology biological process included only terms with *P*<0.001 and with at least 3 enriched genes.

### UMI-4C analysis

Paired-end reads were demultiplexed according to the viewpoint sequence using fastx-multx from ea-utils^57^ and analyzed using umi4cPackage^27^. 4C tracks were created by selecting viewpoint-specific reads, aligning them to the genome and extracting the number of UMIs using the *p4cCreate4CseqTrack* function (see quality control statistics in **Supplementary Table 9**). Cytokine-treated profiles were then scaled to the control profile using the umi4cPackage package function *p4cSmoothedTrendComp.* Profiles were also smoothen based on the total number of UMIs present in a 2Mb region centered on the viewpoint and excluding 3kb around it. The following formula was used to calculate the minimum UMIs needed for smoothing. If the fragment did not reach this minimum, it was then merged with successive fragments until minimum was reached.

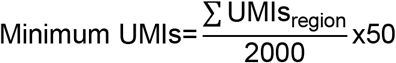

For detecting differential contacts, we focused on a 2Mb region centered on the viewpoint, but excluding 1.5kb on each side of the viewpoint. We then partitioned the region into windows of width proportional to the mean restriction fragment length in the region (mean_fragment_):

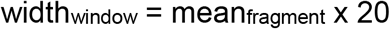

Differential contacts analysis was performed only in those windows containing at least one regulatory element, thus limiting the number of tests, using the *p4cIntervalsMean* function from the umi4cPackage, which uses a Chi-squared test to assess significance. P-values were then corrected for multiple-testing using the FDR method. Windows with an FDR adjusted *P* < 0.05 were considered significant. To quantify the chromatin contact changes, we counted the number of cytokine-treated and control UMIs for each window and computed their odds ratio based on the total UMI counts in the region, following the formula

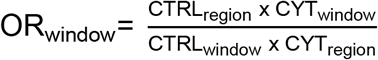

where CTRL and CYT represent the number of UMIs in control (CTRL) and cytokine-exposed (CYT) conditions.

### Risk loci analyses

We first selected from the NHGRI-EBI GWAS Catalog^58^ all “leading” SNPs with disease trait matching either “Type 1 diabetes” or “Type 2 diabetes” (date 14-06-2018). Next, we extended our collection of associated variants to all those in strong LD (R^2^>0.5, CEU) with the lead SNP (source of LD information 1000 Genomes Project phase3^59^). Disease-associated loci boundaries were defined by setting the coordinates of the first and the last SNP in LD at each locus, overlapping regions were merged. In order to focus on unambiguous traits, T1D and T2D shared risk loci were removed for this analysis.

To calculate if the overlap of IRE or SRE enhancers with disease-associated loci exceeded random expectations, we performed permutation tests using the R package regioneR^50^. All randomizations were restricted to mappable genome coordinates by using a masked version of hg19 genome and risk loci were permuted 5,000 times for each comparison. Significance was established at *P*<0.05.

Leading SNPs and their proxies (LD>0.5) directly overlapping IREs in human islet samples were annotated using *findüverlaps* function from GenomicRanges R package^60^. In **Supplementary Table 3** we show a comprehensive dataset of T1D risk loci containing at least one IRE consistently induced in both cell types (as defined above) or one human islets IRE.

### GWAS association analysis

Genome wide association statistics were generated from 5,909 type 1 diabetes cases (from the UK GRID^61^ cohort) and 8,721 controls (1,397 from the UK blood service^62^, 5,472 from the 1958 British birth cohort^63^ and 1,852 from the bipolar disorder cases from the welcome trust case control consortium^62^). 6,653 (1,926 cases and 4,727 controls) were genotyped using the Affymetrix GeneChip 500k array and 7,992 (3,983 cases and 3,994 controls) were genotyped using the Illumina Infinium 550k array. Prior to imputation, we filtered SNPs with a call rate of less than 95%. Shapeit2^64^ was used to prephase individual haplotypes using the haplotype reference consortium (HRC) as the haplotype reference panel, then minimac3^65^ was used to impute both batches genome-wide for chromosomes 1-22. As post imputation quality control, we excluded SNPs with a Minor Allele Frequency (MAF)<0.01 (or MAF<0.05 if the SNP was only present in one of the genotyping batches), an imputation information score <0.3 a difference in MAF between Illumina and Affymetrix controls of >5%, a difference between Illumina and Affymetrix cases of >5% and a difference in MAF of >5% between combined controls and European members of the HRC. As a further quality control (QC), we removed SNPs where the difference in log-odds ratio between cases and controls was >0.3, or the odds ratio difference was >0.3 and the effects were in opposite directions between batches. Finally, if a SNP was present in only one of the genotyping batches, we removed SNPs with an imputation r^2^ of less than 0.8. This conservative QC procedure was implemented to minimize the probability of observing spurious associations due to imputation.

SNPTEST^66^ was used to test the association of each SNP with type 1 diabetes status, adjusting for the top 3 principal components, for those genotyped with the Illumina chip and the Affymetrix chips separately, then results were combined in an inverse-variance weighted fixed effects meta-analysis.

Once association results were generated, we filtered-out rare variants with MAF < 0.01. For the definition of 99% credible sets we followed previously defined methodology^67^. Shortly, we took all SNPs with *P*<5e^-8^ and created a 1Mb window around them, merging all overlapping windows. This allows us to generate the potential risk loci; we considered as the top SNP of the locus the one with the smallest P-value. Next, taking advantage from the R^2^ scores from 1,000 Genomes phase 3 we removed from further analysis those variants whose association (R^2^) with the top SNP in the locus was < 0.1. Finally, we calculated the Approximate Bayes Factor (ABF) and Posterior Probability (PP) of each variant within the locus, and included in the credible set all those variants until reaching a cumulative PP over 0.99.

**Supplementary figure 1.**
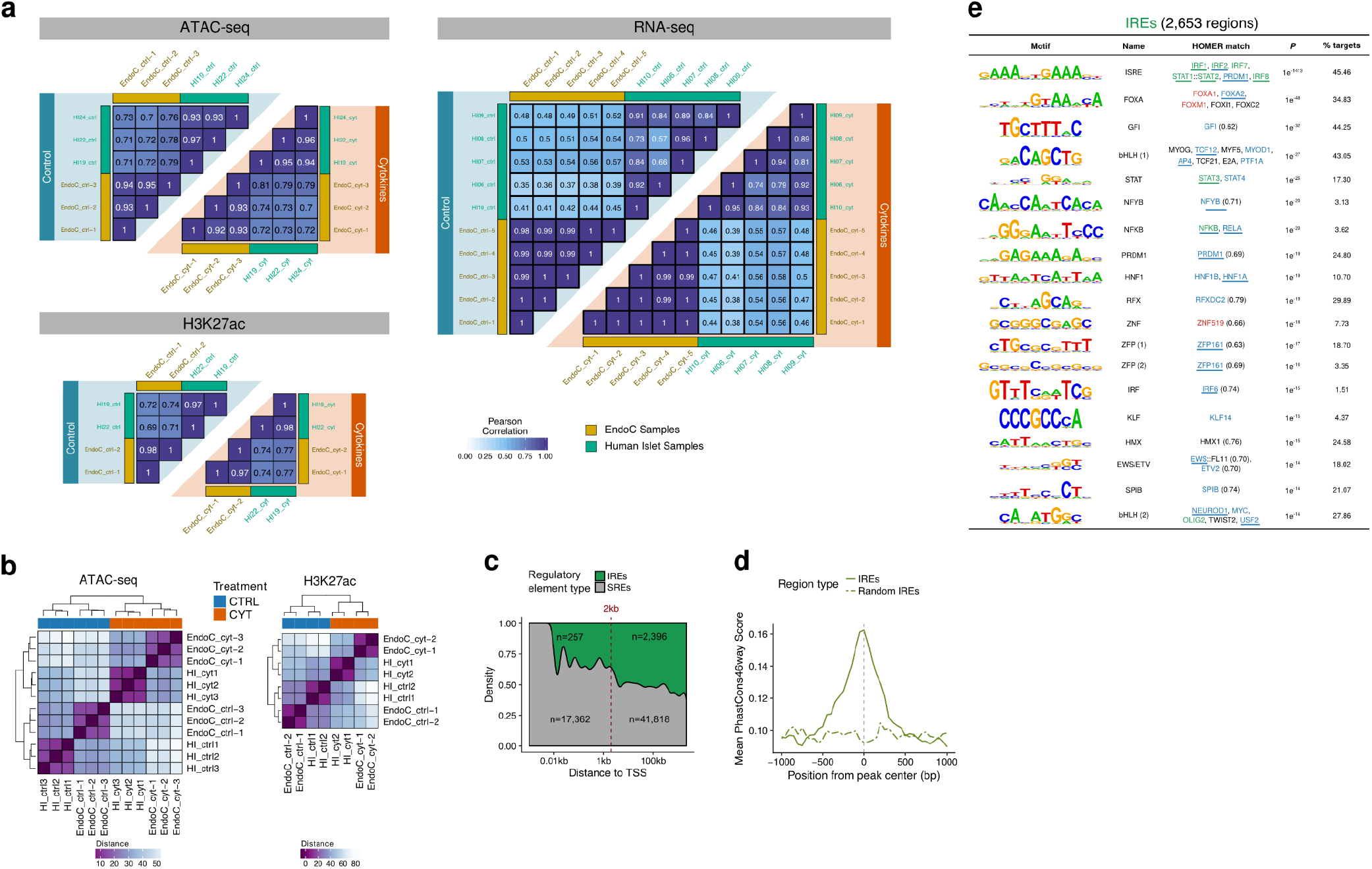
Chromatin and transcriptomic characterization of human pancreatic β cells exposed to pro-inflammatory cytokines. **a**, Pearson correlation between replicates in the different assays and conditions. High correlation levels are observed within replicates of the EndoC-βH1 β cell line and within biological replicates of human pancreatic islets primary tissues (HI 1-9). Importantly, we still observe high correlation when comparing human islets and EndoC-βH1. Lower correlation values in the latter comparison, compared to inter-replicates correlations are due to the heterogeneity of the human islet cell population composed primarily of β-cells (~50-60%) but including other cell types such as α, γ, δ or ε cells. **b**, Hierarchical clustering using normalized ATAC-seq and H3K27ac ChIP-seq read counts at IREs common in EndoC-βH1 and human islets (see Methods) show that samples cluster according to treatment and not to sample type or biological background. Interestingly, IREs allow classifying both ATAC-seq and H3K27ac samples according to the treatment, suggesting that the differences caused by the proinflammatory cytokines are greater than those derived by the sample heterogeneity. HI=Human pancreatic islets, EndoC=EndoC-βH1 **c**, Distribution of distances to nearest TSS for the different types of regulatory elements, showing that IREs, compared with stable regulatory elements (SREs), are preferentially located distally to TSS. **d**, Mean sequence conservation score of IREs and a randomized set of IREs in placental mammals. Peaks were extended from the center 1kb to each direction and mean score was calculated in 50bp windows. **e**, Sequence composition analysis of IREs (n=2,653) illustrating the top identified *de novo* motifs. Colors for matched genes correspond to RNA-seq (name) or protein (underlined) status (red=down-regulated, blue=equal-regulated, green=up-regulated, black/no line = not expressed/detected).

**Supplementary figure 2.**
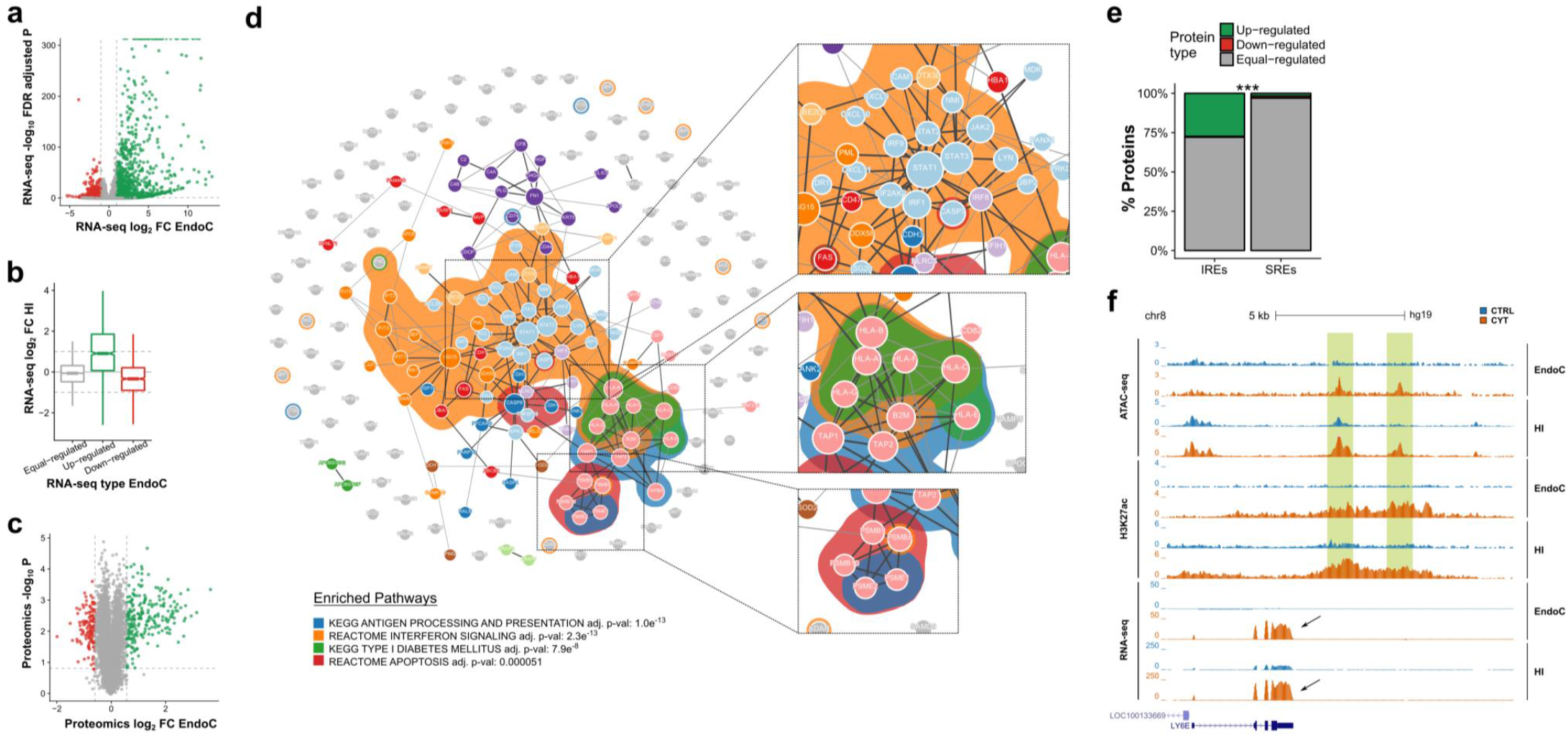
Exposure to pro-inflammatory cytokines drives changes in the transcriptome and proteome of pancreatic β cells. **a**, Volcano plot of RNA-seq genes, showing up-regulated genes (green) and down-regulated genes (red) upon exposure of EndoC-βH1 to cytokines. Vertical lines indicate the log_2_ FC threshold (|log_2_ FC|>1) and horizontal line indicates the FDR adjusted *P* cutoff for significance (FDR adjusted *P*>0.05) calculated by fitting a negative binomial model in DESeq2. **b,** Distribution of RNA-seq counts in human islet samples in the genes previously classified as up, down or equal-regulated in EndoC-βH1 cells. Boxplot limits show upper and lower quartiles, whiskers extend to 1.5 times the interquartile range and the notch represents the confidence interval around the median. **c**, Volcano plot for multiplex proteomics, showing in green the up-regulated proteins and in red the down-regulated, which have a Q-value<0.1 and |log_2_ FC|>0.58. Vertical lines indicate the log_2_ FC thresholds. **d**, Protein-protein Interaction (PPI) network generated from up-regulated proteins after cytokine exposure. Node color indicates belonging to same interacting community and background corresponds to specific pathway enrichment. **e**, Proportion of up, equal or down-regulated proteins encoded by genes located <20 kb from IREs or SREs. *** *P*<0.001 Chi-squared test. **f**, View of the *LY6E* locus, whose expression is induced after cytokine exposure and coupled with chromatin changes in the vicinity.

**Supplementary figure 3.**
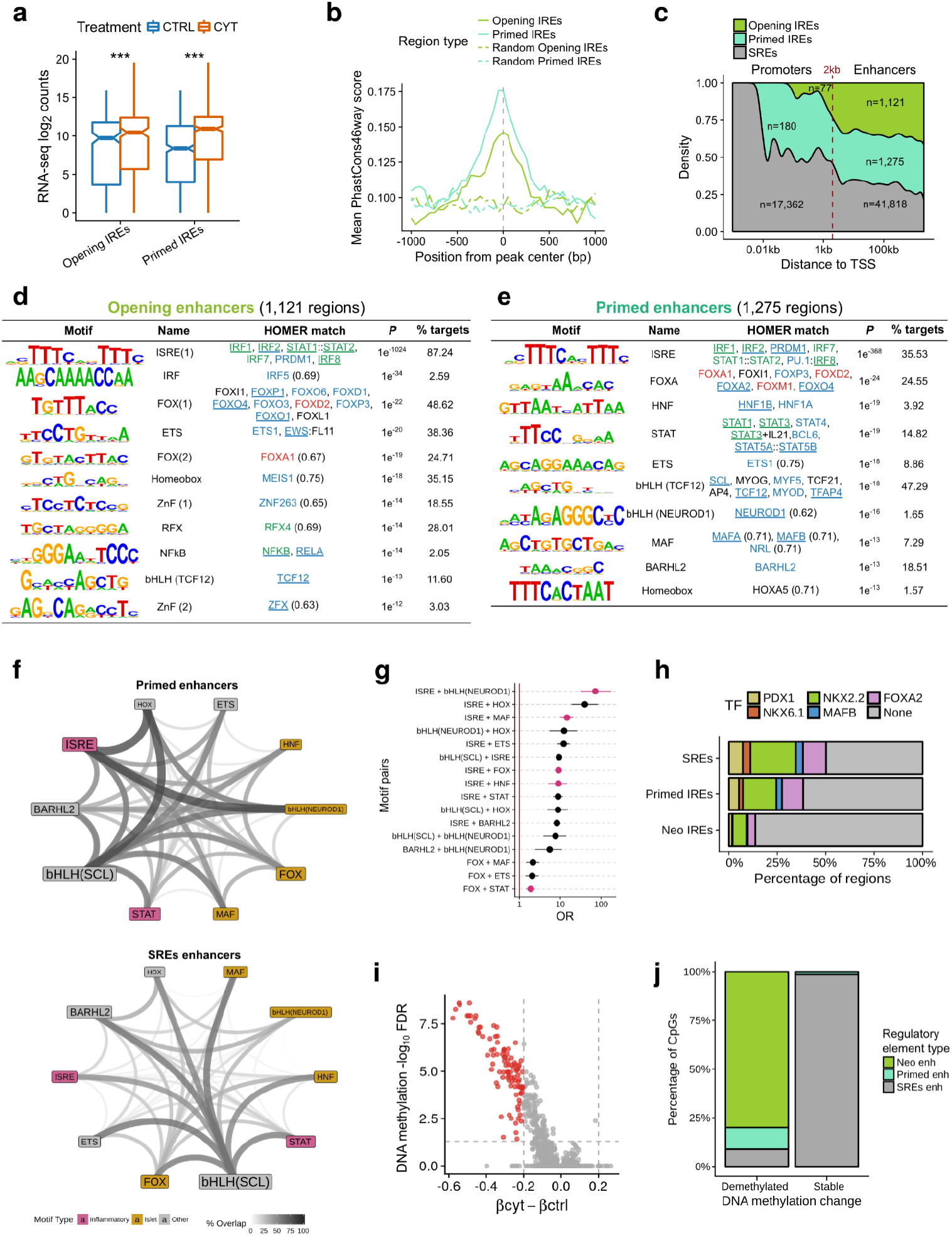
Characterization of β cell IREs. **a**, Genes located <20 kb from different classes of IREs (see **Fig 3a** for the classification) show cytokine-induced expression in EndoC-βH1 cells exposed or not to pro-inflammatory treatment. CYT= cytokine exposed (orange), CTRL = not exposed to cytokine (blue). Boxplot limits show upper and lower quartiles, whiskers extend to 1.5 times the interquartile range and the notch represents the confidence interval around the median. *** *P*<0.001 Wilcoxon test. **b**, Mean sequence conservation score of the two classes of IREs (solid lines) and a randomized set of each (dotted lines) used as control. Scores were obtained from placental mammals, the peaks were extended 1kb to each direction from the center and mean score was calculated in 50bp windows. **c**, Distribution of distances to nearest TSS of the different classes of open chromatin as indicated in **Fig. 3a**. Dotted line indicates the threshold used in the following analyses to classify them in “promoters” (|distance to TSS|<2kb) or “enhancers”. **d, e**, Top hits for *de novo* motif analysis in opening enhancers (n=1,121) **(d)** and primed enhancers (n=1,275) showing that primed enhancers are enriched in inflammatory response and islets-specific (FOXA2, HNF1A/B, MAFA/B, NEUDOD1) TF binding motifs while opening enhancers include exclusively inflammatory response TF recognition sequences **(e)**. Colors for matched genes correspond to RNA-seq (name) or protein (underlined) status (red=down-regulated, blue=equal-regulated, green=up-regulated, black/no line = not expressed/detected). **f**, Diagram showing the percentage of colocalization between the TF binding sites identified by *de novo* motif analysis in primed enhancers. The colocalization was computed in primed enhancers (top) and, as a comparison, in SRE enhancers (bottom). The size of the label indicates the number of regions containing the TF binding sites, color indicates the type of motif and the width and intensities of the lines indicates the percentage of regions in which two motifs map to the same open chromatin site. **g**, Odds-ratio of the probability of finding a motif pair in the same open chromatin site in primed enhancers compared to SRE enhancers. Only significant pairs (FDR adjusted *P*<0.05) are shown. The analysis presented in **f** and **g** show that immune and islet-specific TF motifs colocalize to the same chromatin site significantly more often in primed enhancers compared to SRE. **h**, Percentage of overlap between EndoC-βH1 SRE, primed and neo enhancers and different islet-specific TFs binding obtained from ChIP-seq assays in untreated human pancreatic islets. **i**, Volcano plot showing differentially methylated sites in EndoC-βH1 cells treated or not with the cytokine cocktail. Vertical dotted lines indicate the threshold for differences in methylation (|β-value difference|>0.20; 20% changes in DNA methylation) and horizontal dotted line indicates the threshold for significance (FDR adjusted *P*<0.05, limma moderated t-test). Differentially demethylated sites are shown in red. **j**, Distribution of demethylated and stable CpGs according to the different classes of open chromatin.

**Supplementary figure 4.**
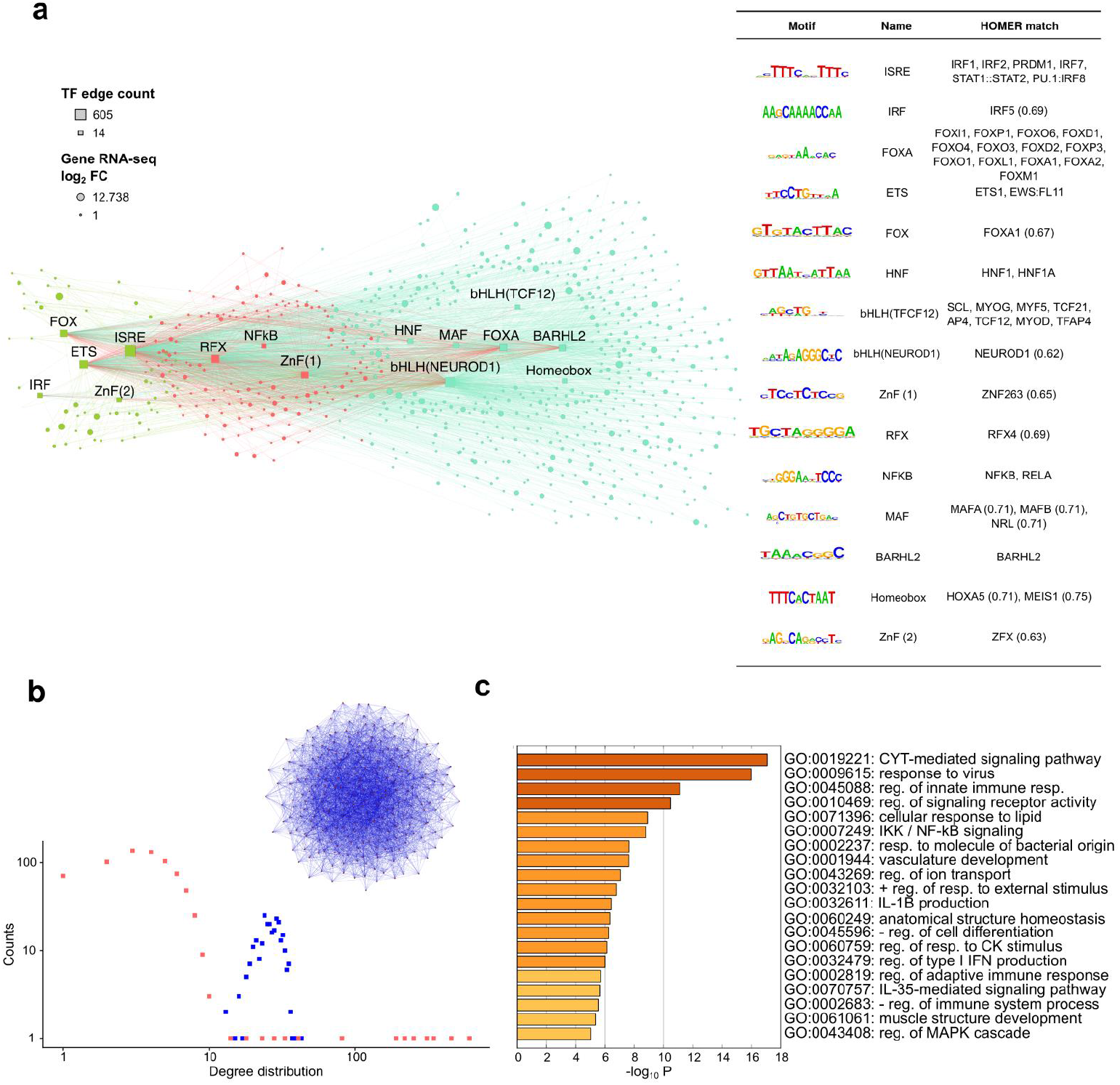
Deconstructing cytokine induced *cis*-regulatory network. **a**, Gene Regulatory Network (GRN) derived from IREs and their putative target genes. Squares represent the IREs inferred TF binding sites (motifs logos and TF matches are shown on the right side) and the ellipses represent their putative target genes (see methods). The squares size reflects the number of connections (edge count) while the gene node size reflects the log_2_ FC of RNA expression after cytokine exposure. The resulting GRN is an interconnected scale-free network composed of 721 nodes and 2,831 edges. Genes regulated exclusively by primed IREs are represented in blue while green depicts opening IREs regulated genes. Red denotes genes regulated by both types of IREs. In each of these three groups the representation of the hierarchy is based on the principle of network centrality where authoritative nodes are located more proximal to the core. **b**, Comparison between the degree distribution of the observed GRN (red) and a random generated network (blue) having the same number of nodes and edges. The bell-shaped degree distribution of random graph denotes a statistically homogeneity in the degree of small and large nodes. In contrast, the observed network showed a high frequency of small degree nodes and a low frequency of highly connected nodes as is typical of a scale-free network. **c**, Bar plot of gene ontology biological process enrichment analysis. Gene-ontology analysis was performed using all target genes in the GRN. Functional enrichment analysis was performed by Metascape (http://metascape.org)^56^ Only terms with *P*<0.001 and with at least 3 enriched genes were considered as significant. Color is proportional to their *P* values.

**Supplementary figure 5.**
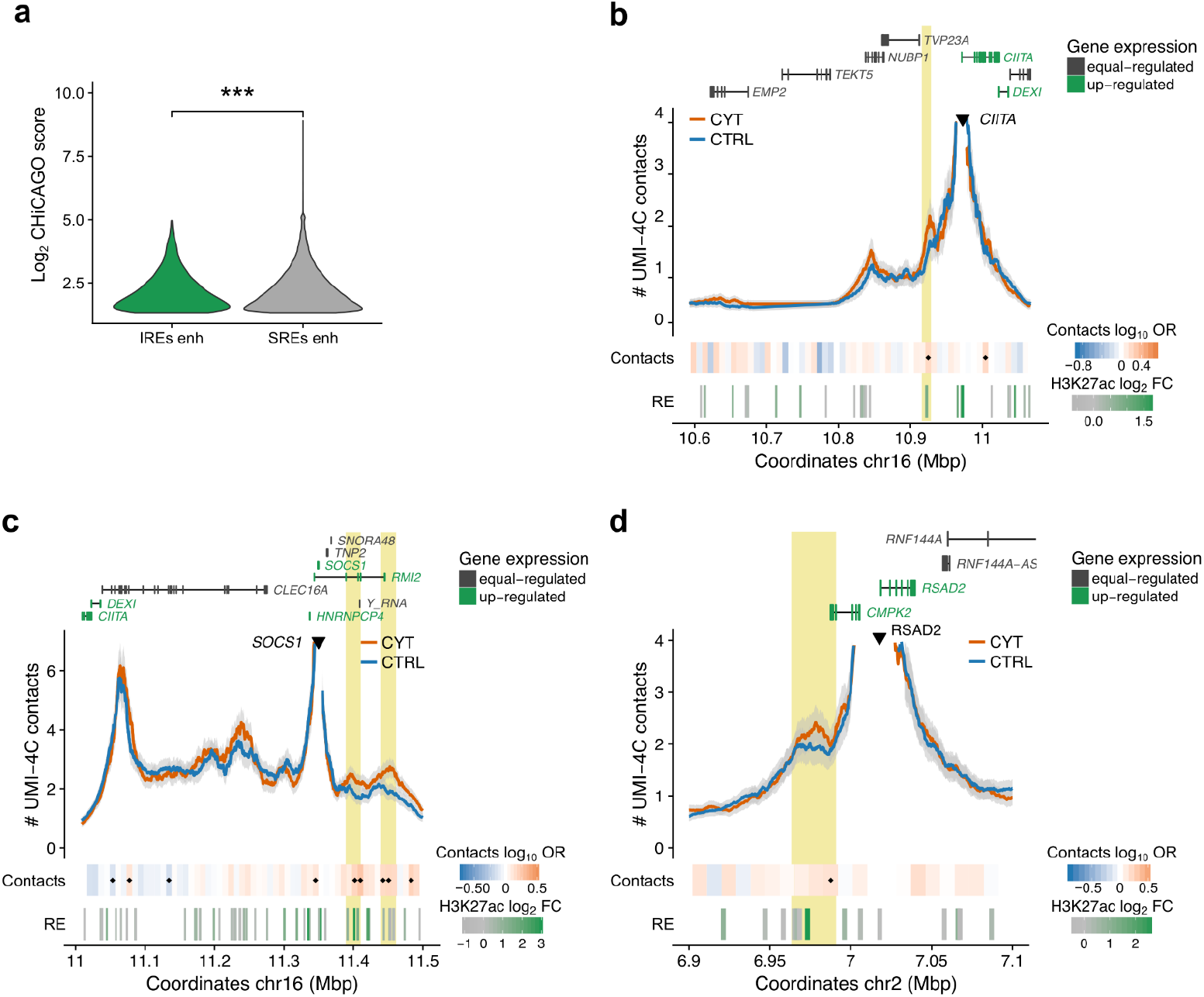
3D chromatin changes induced by exposure of human islets to pro-inflammatory cytokines. **a,** Violin plots showing the distribution of CHiCAGO scores of contacts, detected by pc-HiC experiments in untreated human islets^68^, between different stable and induced enhancers and their target genes. SREs engage chromatin contacts with higher interaction scores compared to those detected for IREs. *** *P*<0.001. **b, c, d**, Views of the 3D chromatin contacts of *CIITA* **(b)**, *SOCS1* **(c)** and *RSAD2* (d) promoters obtained by UMI-4C performed in islets exposed or not to pro-inflammatory cytokines. In yellow we highlight those IREs that gain contacts with the up-regulated gene promoter. A heatmap under the 4C track represents the log_10_ odds ratio (OR) of the UMI-4C contacts difference in cytokine vs. control and a small black diamond on top of the contact heatmap indicates a significant difference in contacts between cytokine-treated and control samples (*P*<0.05). ATAC-seq peaks are represented by rectangles, shaded from gray to green proportionally to the cytokine-induced H2K27ac fold change observed at that site.

**Supplementary figure 6.**
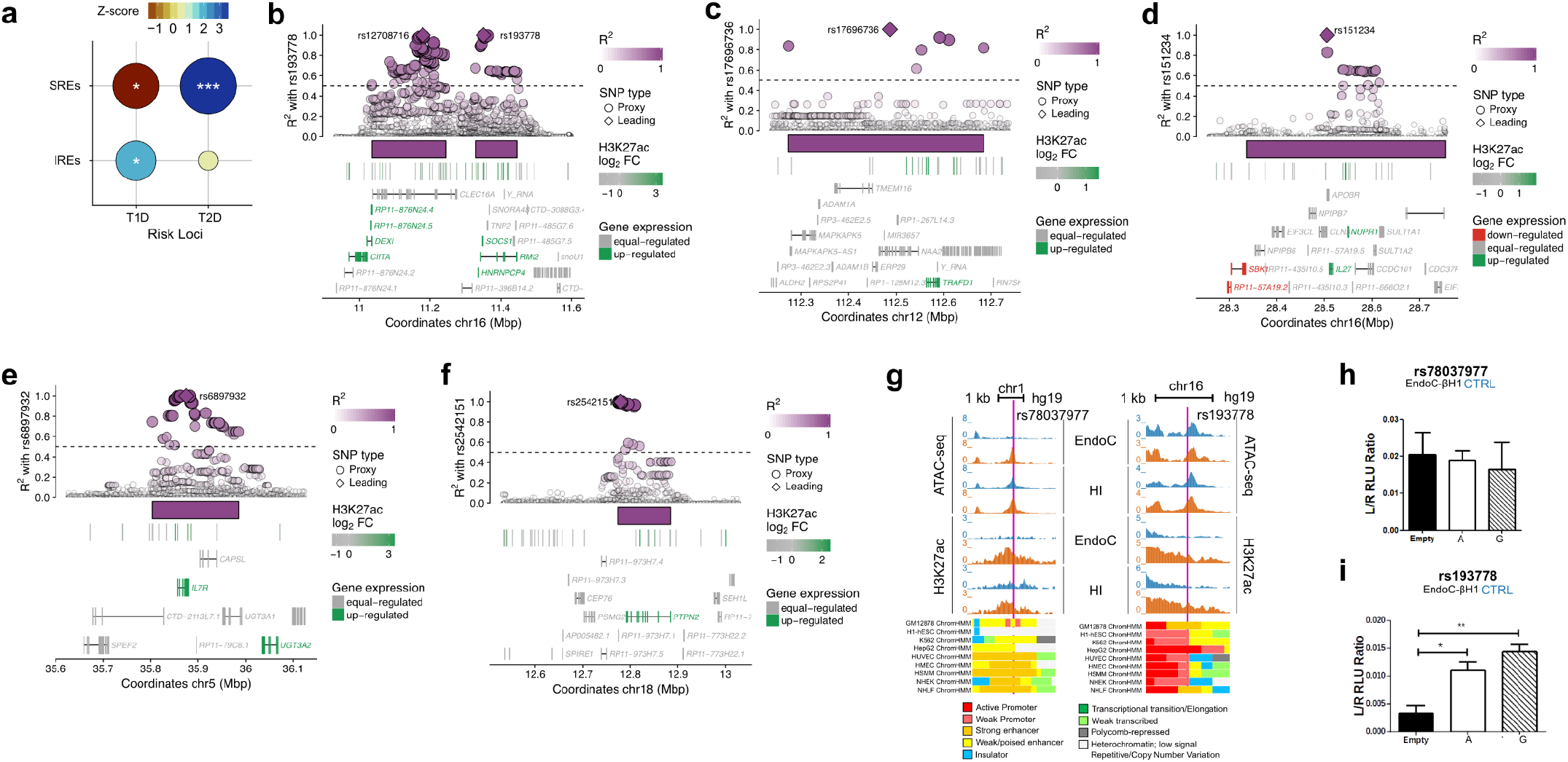
Cytokine-induced islet regulatory elements are enriched in T1D associated loci. **a**, Loci associated with T1D contain EndoC-βH1 cytokine-induced regulatory elements more often than expected while the opposite is true for T2D. The permutation test results assessing the significance of the overlap between types of regulatory elements (y axis) and different trait-associated loci (x axis) are shown. The size of the circle is proportional to −log_10_*P* value; fill represents z-score of the observed versus expected value. Significance assessed by permutation tests; * *P*<0.05, ** *P*<0.01, *** *P*<0.001. **b, c, d, e, f**, Regional plots of different T1D risk loci containing IREs and up-regulated genes. R^2^ values are based on 1KG CEU and the leading SNPs in the locus is represented by a diamond. Dotted line represents the LD cutoff used for this analysis (LD>0.5). **g**, The IRE bearing the T1D associated variant rs78037977 is marked by the ENCODE ChromHMM classification as a “strong enhancer” (orange) in other non β-cell lines (left). ENCODE ChromHMM classification in non β-cell lines for the IRE bearing the T1D associated variant rs193778. **h, i**, Allele-specific luciferase experiments for rs78037977 **(h)** and rs193778 **(i)** in untreated EndoC-βH1.

**Supplementary figure 7.**
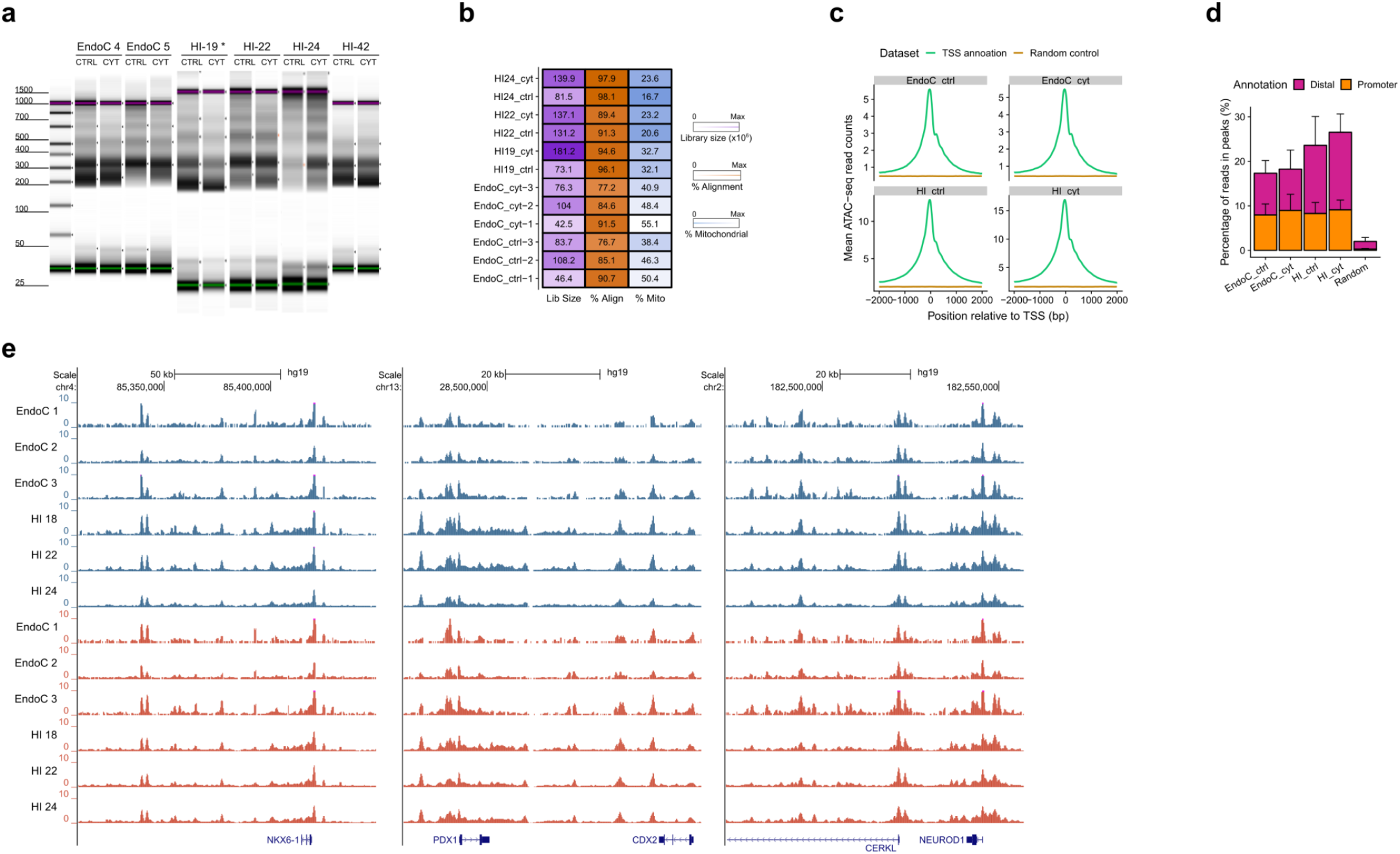
ATAC-seq quality control measures. **a,** Agilent TapeStation profiles obtained by chromatin tagmentation of human islets and EndoC-βH1 samples showing the laddering pattern of ATAC-seq libraries. The band sizes correspond to the expected nucleosomal pattern. *Notice that samples HI-19 CTRL and CYT were used as examples to illustrate the expected fragment distribution pattern in ATAC-seq experiments in Raurell-Vila et al.^12^. **b**, Summary of per-replicate sequencing metrics, showing total library sizes, percentage of aligned reads and percentage of mitochondrial aligned reads. **c**, TSS enrichment over a 4kb window centered on genes TSS compared to a set of genes randomized along the genome, showing the expected pattern of open chromatin centered on the TSS. **d**, Percentage of total reads found at called open chromatin peaks classified as distal (>2kb from TSS) or promoters (≤2kb from TSS) compared to a randomized set of peaks. **e**, UCSC views at islet-specific loci *(NKX6.1, PDX1* and *NEUROD1)* showing the high reproducibility of ATAC-seq profiles among independent replicates.

